# Single nuclei RNAseq analysis of HD mouse models and human brain reveals impaired oligodendrocyte maturation and potential role for thiamine metabolism

**DOI:** 10.1101/2022.06.27.497613

**Authors:** Ryan G. Lim, Osama Al-Dalahmah, Jie Wu, Maxwell P. Gold, Jack C. Reidling, Guomei Tang, Miriam Adam, David Dansu, Hye-Jin Park, Patricia Casaccia, Ricardo Miramontes, Andrea M. Reyes-Ortiz, Alice Lau, Fatima Khan, Fahad Paryani, Alice Tang, Kenneth Ofori, Emily Miyoshi, Neethu Michael, Nicolette Geller, Xena E. Flowers, Jean Paul Vonsattel, Shawn Davidson, Vilas Menon, Vivek Swarup, Ernest Fraenkel, James E. Goldman, Leslie M. Thompson

**Affiliations:** UCI MIND, University of California Irvine, CA; Department of Pathology and Cell Biology, Columbia University Irving Medical Center, New York City, NY; Department of Biological Chemistry, University of California Irvine, CA; Department of Biological Engineering, Massachusetts Institute of Technology, Cambridge, MA; Department of Neurology, Vagelos College of Physicians and Surgeons, Columbia University Irving Medical Center, New York City, NY; Psychiatry and Human Behavior, University of California Irvine, CA; Neurobiology and Behavior, University of California Irvine, CA; Department of Pathology, University of California Irvine, CA; Sue and Bill Gross Stem Cell Center University of California Irvine, CA; Taub Institute for Research on Alzheimer’s Disease and the Aging Brain, Columbia University Irving Medical Center, New York City, NY; Lewis-Sigler Institute for Integrative Genomics, Princeton, NJ; Advanced Science Research Center at the City University of New York

**Author notes:** co-first authors (Ryan G. Lim, Osama Al-Dalahmah, Jie Wu).

## Abstract

The complexity of affected brain regions and cell types is a challenge for Huntington’s disease (HD) treatment. Here we used single nucleus RNA sequencing (snRNAseq) to investigate mechanism of pathology in the cortex and striatum from R6/2 mice at 8 and 12w and in three regions of human HD post-mortem tissue. We identified cell type-specific and cell agnostic signatures and found changes suggesting oligodendrocytes (OLs) and oligodendrocyte precursors (OPCs) were arrested in intermediate maturation states. OL-lineage regulators OLIG1 and OLIG2 were negatively correlated with CAG length in human OPCs, and ATACseq analysis of HD mouse NeuN-negative cells showed decreased accessibility of sites regulated by OL maturation genes. Glucose and lipid metabolism were implicated in abnormal cell maturation and PRKCE and *Thiamine Pyrophosphokinase 1* were identified as central genes. High dose thiamine/biotin treatment of R6/1 HD mice to target thiamine metabolism not only restored OL maturation, but also rescued pathology in neurons. These findings reveal insights into HD OL pathology that spans multiple brain regions and link OL maturation deficits to abnormal thiamine metabolism.

## Introduction

Huntington disease (HD) is a progressive neurodegenerative disease characterized by prominent loss of medium spiny neurons (MSN) in the striatum and cortical atrophy ^1^. The disease, which manifests with cognitive, psychiatric and movement impairments, is caused by an autosomal dominant CAG repeat expansion in the first coding exon of the Huntingtin gene and a corresponding expanded polyglutamine repeat in the Huntingtin (HTT) protein ^2^. Genome-wide approaches, including bulk RNA- and ChIP-sequencing, have facilitated understanding the molecular impact of mutant HTT (mHTT) expression in a variety of model systems ^3–6^ and have suggested deficits in neurodevelopmental programs in HD ^3,7–9^, however bulk tissue analysis limits understanding of cell type-specific changes. The ability to distinguish common signatures of HD across multiple cell types from those unique to specific cell types facilitates our mechanistic understanding of disease. Expression of mHTT using cell type-specific drivers in animal models of HD ^10^ or human HD induced pluripotent stem cells differentiated to specific cell types support the idea that cell type-specific effects of HD synergistically lead to pathogenesis ^11,12^. Further, single cell transcriptomics approaches have supported the concept of cell type specific neurodevelopmental impairments in HD.^13,14^

There has been a growing awareness that OL-lineage cells are abnormal in HD. First, early myelination deficits based on structural and transcriptomic studies were described in mouse models of HD^15,16^. OL targeted mHTT expression causes HD symptoms in mice, as well as myelination deficits and altered OL maturation via a mechanism involving Myrf^17^. Myelination deficits due to mHTT expression were evident in spinal cord white matter in BACHD mice^18^. Consistently, bulk transcriptional studies of human HD revealed that *MYT1L*, a myelin transcription factor, and *MBP* were decreased in the caudate and prefrontal cortex, respectively ^19,20^. Second, glial dysfunction^21,22^ and impaired OPC differentiation has been described for HD. For example, HD embryonic stem cell-derived glial progenitors transplanted into shiverer mice exhibit decreased differentiation and hypomyelination compared to controls ^23^. Another study showed that remyelination was impaired in cuprizone-treated mice, implicating abnormal OPC function in HD^24^, and inactivation of mHTT in OPCs prevented myelin abnormalities in HD mice ^11^. Clinical radiographic and neuropathological studies also reveal that OLs and myelination are abnormal in human HD (summarized in ^25^). Neuropathologic examination of postmortem HD brains revealed higher density of OLs in the caudate nucleus^26 27^, including in pre-symptomatic HD patients. Stereological examinations of white matter reveal a decrease of 20-30% of the cross-sectional area of white matter in coronal levels from frontal to occipital regions ^28^, as well as in the fornix^29^, in both lower and higher HD grades, suggesting that white matter loss represents an early change.

Here, we used single nucleus-RNAseq (snRNAseq) to obtain cell type-specific gene expression data across multiple brain regions from both the rapidly progressing R6/2 mouse model ^30^ and human post-mortem brain samples with increasing grades of disease severity – including both adult- and juvenile-onset HD - and used these data for correlative and causal network modeling. We identified cell type-specific and agnostic gene expression changes, as well as putative causal drivers of transcriptomic changes. Consistent with previous literature, we find that oligodendrocyte-lineage cells show significant transcriptional dysregulation. Expanding on these findings, HD OPCs and OLs have altered expression of development and maturation genes in both mice and human tissue, with many HD OL-lineage cells showing intermediate states of development. The extent of dysregulation correlates with CAG repeat length in human tissue; the same dysregulated genes were also highlighted by causal modeling in our mouse data. A gene central to the OPC/OL causal network, Protein kinase C epsilon (PRKCE), was downregulated in human and mouse tissue, and functional studies clarified its role in promoting OL maturation. Evidence from ATACseq and validation studies support this dysregulation. Notably, we identify impairments in glucose and lipid metabolism, identified as cell type agnostic signatures, as potential drivers of this pathology. This connection to metabolism led us to find potentially unique roles for diacylglycerol (DAG), and thiamine and biotin (T&B) metabolic processes in HD OL maturation impairments. Thiamine Pyrophosphokinase 1 (*Tpk1)*, which converts thiamine into thiamine pyrophosphate, was differentially expressed in the most cell types in the 12w R6/2 mice, and both *TPK1* and *SLC19A2*, a thiamine transporter, were downregulated in the human HD snRNAseq data. Mutations in TPK1 or the thiamine-transporters SLC19A3 lead to thiamine pyrophosphate deficiencies and early-onset neurodegeneration with brain atrophy, basal ganglia impairment, and motor dysfunction which can be effectively treated with high dose thiamine and biotin (T&B) ^31,32^. In addition, mutations in SLC19A2 lead to Roger’s syndrome, with megaloblastic anemia, thrombocytopenia, diabetes mellites, and sensorineural deafness ^33^ and general dietary thiamine deficiencies are known to contribute to a number of neurological and psychiatric symptoms ^34^. To further examine potential connections between metabolic changes in HD and OL maturation we treated R6/1 mice, which has a longer therapeutic window than R6/2 mice and also show dys-maturation signatures in a number of cell types ^14^, with T&B and conducted snRNAseq on the striatum of T&B treated and vehicle treated mice. T&B treatment resulted in significant rescue of dys-maturation signatures in OL and neurons, and an overall decrease in the number of significant differentially expressed genes (DEGs). Our novel data provide evidence that dysregulated metabolism and metabolic genes can directly contribute to the cell maturation deficits observed in OLs and other cell types, and that diet supplementation may be a therapeutic modality for HD.

## Results

### Single nuclei RNAseq of R6/2 mouse model of HD

R6/2 mice are a rapidly progressing transgenic mouse model that express mHTT exon 1 and have features in common with human symptomatic HD, including transcriptional changes ^30^. To uncover progressive, cell type-specific, and region-specific transcriptional changes, snRNAseq was conducted on three striatal and cortical samples each from R6/2 and non-transgenic (NT) mice at 8w and 12w of age (**Fig. 1a**, See Methods. snRNAseq data were also generated and analyzed from human HD and control brains (**Fig. 1a** **and** **e**, described below). Initial QC and filtering led to the identification of 108,974 nuclei in total. **Fig. 1b** and **Supplementary Fig. 1a** show uniform manifold approximation and projection (UMAP) plots of these data. Unsupervised clustering identified 13 clusters in the 8w and 12w striatal samples, and 18 and 16 clusters in the 8w and 12w cortical samples, respectively (**Fig. 1b**). A select number of cell type gene markers used to annotate these clusters is shown in **Supplementary Fig. 1b**. R6/2 and NT cells clearly separate in some of the clusters. For example, 12w D1+ MSNs completely separated into distinct clusters, which is reflected by the large number of DEGs between the two conditions (**Fig. 1b-d, Supplementary Table 1**). The proportion of cells in each cluster across the cortex and striatum is shown in **Fig. 1d**. We also find large numbers of DEGs in the excitatory (Ex) and inhibitory (Inhib) neurons, astrocyte (Astro), OLs, and OPC clusters (**Fig. 1c**). Minimal to no changes were seen in the microglia (MG), vascular cells, and cholinergic neurons (**Fig. 1c**). These clusters had the smallest number of cells and therefore could lack the power required to identify statistical differences. Regional differences are reflected by differences in cell type-specific DEGs across regions (**Fig. 1c**). The total numbers of DEGs across all cell types were compiled and compared between 8w and 12w samples showing a large overlap of DEGs, with more unique DEGs in the 12w samples for both the striatum and cortex (**Supplementary Fig. 1c**). When we combined all data from both ages and regions, we found no clustering differences for each cell type between age and region, except for cell types that were specific to either the striatum or cortex, e.g. MSNs in the striatum (**Supplementary Fig. 1d**). The only differences between the age groups were seen between the 8w and 12w OLs.

**Fig. 1.**
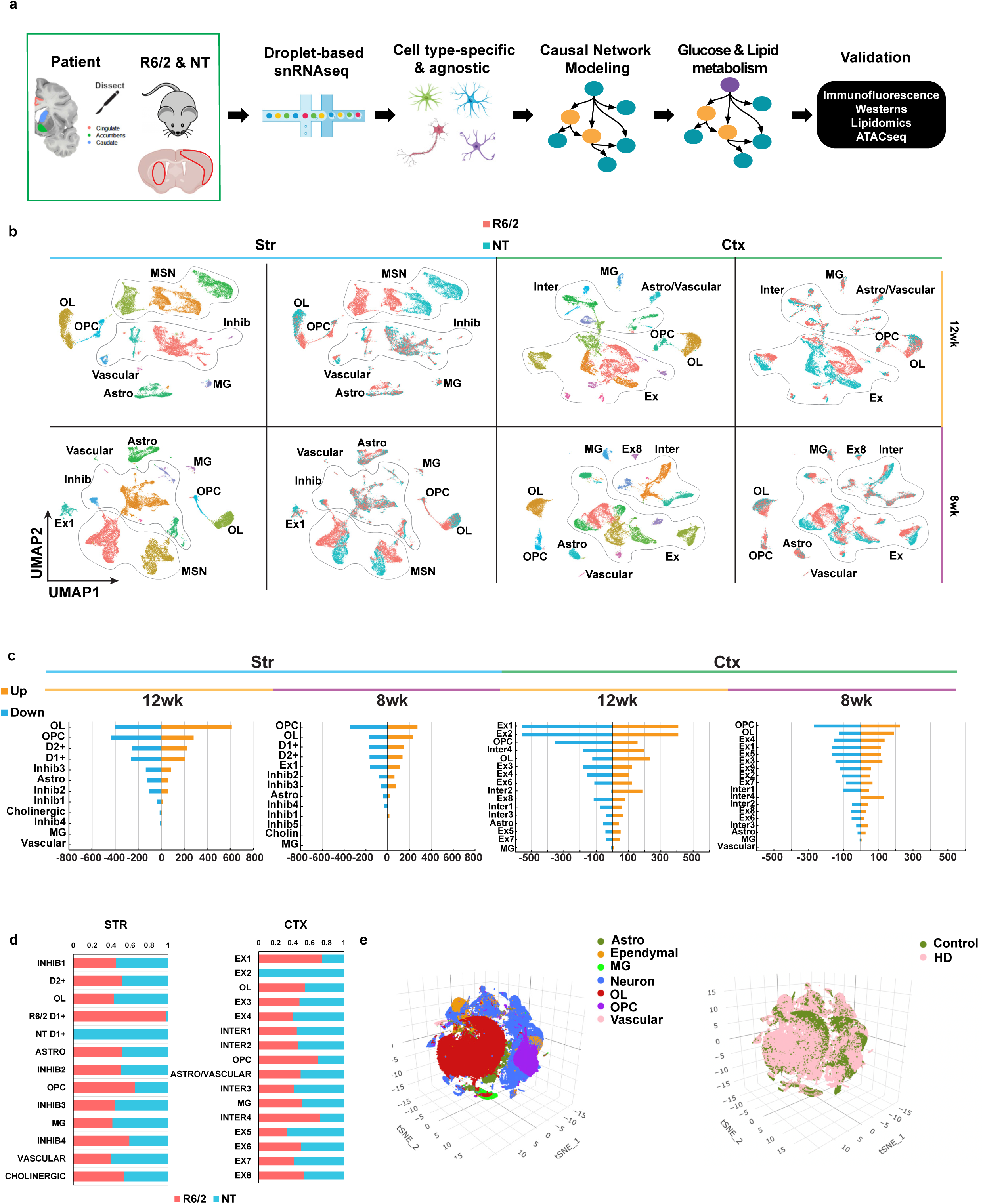
Single nucleus RNAseq of mouse and human R6/2 and HD samples. **a**) Illustration of workflow used for this study. After frozen tissue is microdissected from the Cingulate, Caudate, and nucleus Accumbens from 66 samples from 29 human donors (3 grade I, 4 grade II, 4 grade III, 3 grade IV, 5 juvenile-onset HD, and 10 matched controls), or the striatum and cortex of the mice (*n* = 3), nuclei are isolated, 10X Libraries are prepared followed by next generation sequencing. **b**) Uniform manifold projection and approximation plots (UMAP) of the R6/2 and NT mouse data colored by cluster or genotype. General cell type annotations: Astro = Astrocytes, OL = Oligodendrocyte, OPC = Oligodendrocyte progenitors, MSN = Medium spiny neurons, Inhib = inhibitory neurons, MG = Microglia, Ex = Excitatory neurons, Inter = Interneurons. **c**) Barplot showing the number of up (blue) and down (orange) regulated DEGs per a cell type in the mouse data. **b** and **c**) Striatal (Str, light blue bar) samples on the **left** and cortical (Ctx, light green bar) samples on the **right**, 12w samples marked by yellow bar and 8w marked by purple bar. **d**) Proportion of R6/2 and NT cells within each cluster, red = R6/2 & blue = NT. **e**) tSNE plots of the human snRNAseq results showing color-coded by cell type (**Left**), condition (**Right**), anatomic region (**Bottom Left**), and grade (**Bottom Right**). Right, dotplot showing expression of cell type markers per cluster.

Gene ontology (GO ^35^) enrichment analysis and KEGG pathway analysis were used to investigate the biological implications of each set of DEGs from the different cell types. The top 10 significant terms revealed that the majority of DEGs, regardless of cell type, are involved in neuronal related functions, including neurogenesis, synaptic function, and glutamate related signaling (**Supplementary Fig. 2a**). Certain cell types were enriched for terms such as “developmental process” in OLs and OPCs. Similar to GO analysis across regions, age, and cell type, there were recurring KEGG pathways as well as sets of unique pathways that group together to indicate functional impairment such as focal adhesion, cytoskeleton, ErbB and axon guidance as the top pathways in OLs, suggesting a loss of signaling pathways involved in cell-to-cell communication between OLs and neurons (**Supplementary Fig. 2b**). We also identified cell type agnostic DEGs that were common to both glia and neurons. **Fig. 2a** **and Supplementary Fig. 3a** show the top multi-cluster DEGs identified in at least 50% of the cell types/clusters per tissue region and age, as a heatmap with hierarchical clustering. Many DEGs across both glia and neurons are involved in RNA processing and splicing and metabolism. Hierarchical clustering shows grouping of genes with similar functions indicating potential correlated expression and regulation. KEGG pathway analysis also highlighted metabolic pathways including TCA cycle, O-glycan biosynthesis, amino and nucleotide sugar, sucrose, and pentose phosphate pathways, many of which appear in the earlier 8w age stage (**Supplementary Fig. 2b**). Dysregulated metabolic genes were found in or downstream of the glucose super metabolism pathway that includes glycolysis, the hexosamine biosynthetic, polyol, and diacylglycerol pathways. The two genes dysregulated across the most cell types in the 12w striatum were *Tpk1*, and *Malat1,* a long non-coding RNA involved in RNA processing and transcriptional dysregulation ^36^ (**Fig 2a**). Moreover, *Tpk1* was also among the top dysregulated genes in the 12w cortex, and another glycolytic gene, glucose-6-phosphate isomerase 1 (*Gpi1*), was one of the top multi-cluster DEGs in both 8w striatum and cortex (**Fig. 2a** **and Supplementary Fig. 3a)**. Both metabolic genes are upregulated in R6/2. We investigated whether there was an enrichment for KEGG metabolic genes in the DEGs and which metabolic pathways were most impacted; a composite is shown in **Fig. 2b** **(12w striatum) and Supplementary Fig. 3b**. *Tpk1, Ogt, Dgkx genes, and Galnt13*, found in sub-pathways related to glucose and lipid metabolism, are among the most commonly dysregulated genes in all cell types.

**Fig. 2.**
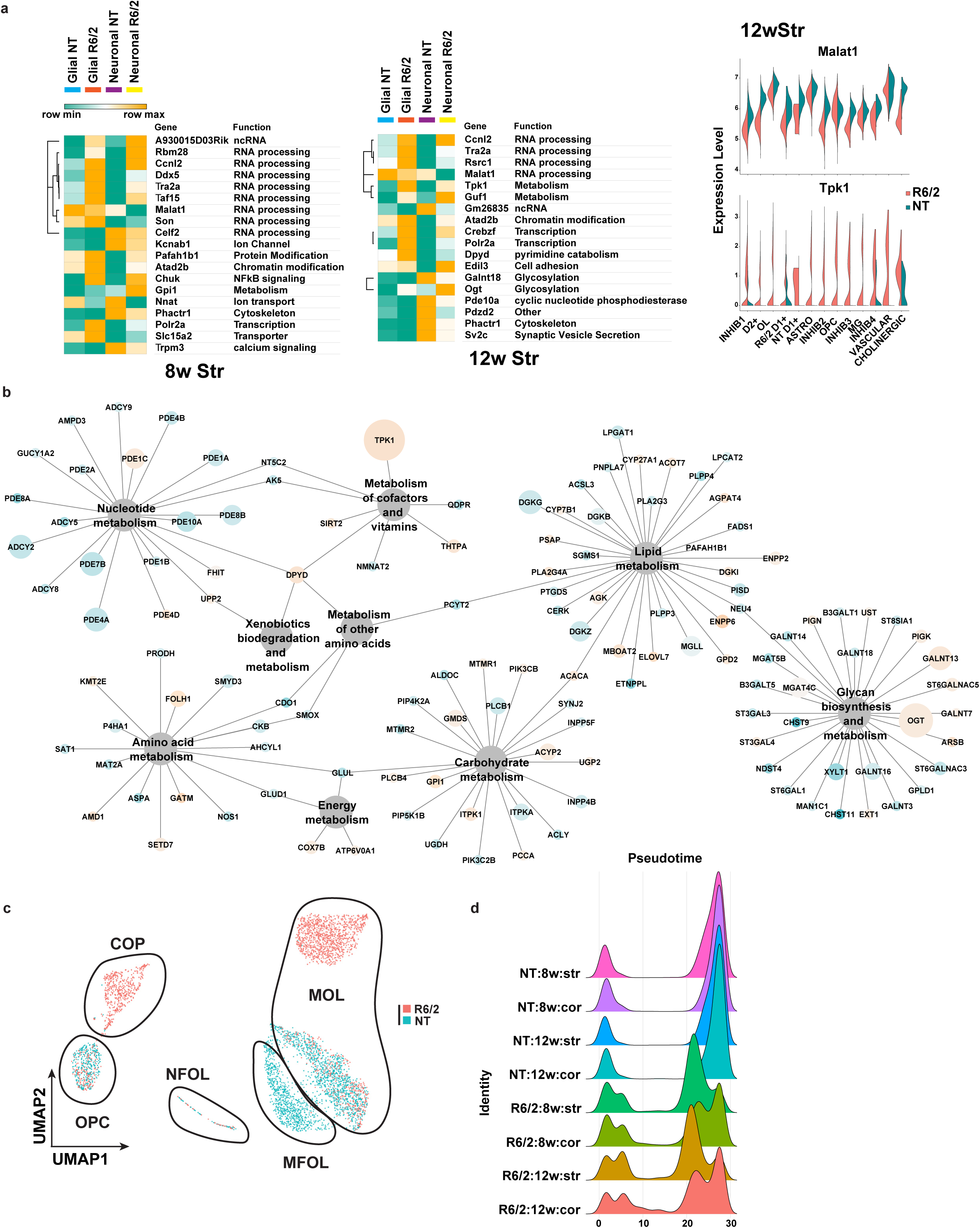
Analysis of differentially expressed genes in R6/2 mice and subclustered analysis of OPCs and OL. **a**) **Left:** Heatmaps and hierarchical clustering of normalized mean expression values in all glial or neuronal cells of the top cell type agnostic DEGs. Cell color represents row min (seafoam green) and max (orange). Color bars denote NT glial cells (light blue), R6/2 glial cells (orange), NT neural cells (purple), and R6/2 neuronal cells (yellow). RNA processing and splicing (*Ccnl2, Tra2a, ddx5, Celf2* and *Taf15*) and metabolism (*Guf1, Tpk1*, and *Gpi1*) related genes. Glucose super metabolism pathway genes that include glycolysis, the hexosamine biosynthetic pathway, polyol pathway, and diacylglycerol pathways, include *Ogt, Tpk1, Gpi1*, and *Galant18*. 8w and 12w Str data shown, cortical data in **Supplementary Fig. 3a. Right:** violin plot of two exemplary genes *Malat1* (**top**) and *Tpk1* (**bottom**) that show global up or down regulation in R6/2 mice, across all cell type, respectively from 12wStr. **b**) Network showing all KEGG metabolic genes significantly dysregulated across the 12wStr DEGs from every cell type. 12w Str data shown, 8w Str and cortical data in **Supplementary Fig. 3b.** Node size is equal to the number of cell types in which the gene is found to be significantly dysregulated and node are colored by up and down regulation (orange = up and blue = down). **c**) UMAPs of subclustered OPCs and OL in the 12w striatum, colored by genotype. Cluster composition: NT cells are mainly MOLs and MFOLs, or OPCs; while R6/2 cells are COP, NFOL, and MOL. Statistical contrasts: R6/2 vs NT for each cluster, cluster comparisons between R6/2 and NT MOLs, NT MFOLs and R6/2 MOL, COP vs OPCs. 8wStr and cortical data show in **Supplementary Fig. 3c. e**) Density plots of cell numbers across pseudotime cell stages, colored by genotype and age.

### R6/2 OPCs are committed to maturation while OLs appear transcriptionally less mature than NT OLs

Given the large changes in OPC and OL clusters, and the UMAPs in **Fig. 1** showing a trajectory of R6/2 cells embedding between the OPC and OL clusters, we investigated whether these cells might represent intermediate cell states between OPCs and OLs. The OL-OPC data were subclustered, revealing six clusters in the 12w striatum and five clusters in the 8w striatum, 8w cortex, and 12w cortex. Each cluster represented distinct populations of OPCs or OLs comprised of R6/2 and/or NT (**Fig. 2c** **(12wk striatum), and Supplementary Fig. 3c-e, integrated data cross regions and ages are described in supplementary results and supplementary Table 2).** These subclustered data were then further annotated based on the gene expression markers and annotations defined by Marques and Zeisel et al ^37^ as OPCs, committed oligodendrocyte precursors (COP), newly formed oligodendrocytes (NFOL), myelin-forming oligodendrocytes (MFOL), or mature oligodendrocytes (MOL) (**Fig. 2c** **and Supplementary Fig. S3c**). DEGs were generated for R6/2 versus NT statistical contrasts for each of the developmental stages. These analyses revealed that R6/2 OPCs (OPC & COP) and OLs (NFOL, MFOL, and MOL) at both ages and in both anatomic regions have changes in expression that suggest developmental/maturation impairments. DEGs included: *Mog, Mag, Mbp, Opalin*, microtubule genes, and genes involved in OL maturation, function, and myelination (**Supplementary Table 1 & Supplementary Fig. 3e**). DEGs involved in glucose and lipid metabolism were also found in OPCs and OLs, including upregulation of *Tpk1*. Pseudotime analysis ^38^ revealed most R6/2 cells were in transitional cell states between OPCs (pseudotime 0) and MOLs (pseudotime 30+), with many HD cells found in the COP cluster and a cluster of NFOL, while NT cells were mostly either OPCs, MFOL, or MOLs (**Fig. 2c** **&** **d** **and Supplementary Fig. 3c-f, these results are further described in the supplement**). HD OL and OPC showed a bimodal distribution at the OPC and OL stages across all ages and regions examined, suggested states of intermediate maturation in both OPCs and OLs (**Fig 2d**). Overall, these data suggest that OPC maturation and subsequent OL differentiation is impaired in R6/2 mice.

### Causal network modeling (CNM) identifies disrupted gene expression networks in R6/2 mice and reveals potential cell type-specific mechanisms of transcriptional change

To investigate disruptions in cell type-specific gene networks in HD, and identify potential key driver genes, we utilized weighted gene co-expression network analysis (WGCNA ^39^) and Bayesian causal network modeling (**Fig. 1a**) to identify causal relationships between genes identified as cell type-specific DEGs and correlated gene network modules ^40–42^. After feature selection (Methods), we used WGCNA and ran a signed network analysis using cells from all NT samples; 6 gene co-expression modules were detected across cortical and striatal tissues at both ages (**Fig. 3a, Supplementary Table 3, and Supplementary Figure 4**). Trait-module correlation analyses showed that our modules were correlated to specific cell types (**Fig. 3a**). The yellow module positively correlated with neuronal cell types and negatively correlated with glia, and the red, turquoise, green, brown, and blue modules positively correlated with Ex, MSNs, MG, Astros, and OLs, respectively. GO enrichment analysis of gene module members showed enrichment for terms related to each cell type (**Fig. 3b**). For example, the OL-correlated blue module was enriched for myelination-related terms. Except for the green module, each module was significantly enriched for DEGs determined using the hypergeometric test (**Supplementary Fig. 5a**), suggesting that these gene networks are relevant to the disease state and become impacted as the disease progresses. The connectivity of the top module members rank-ordered by eigengene-based connectivity (kME) revealed significant alterations (**Fig. 3c**).

**Fig. 3.**
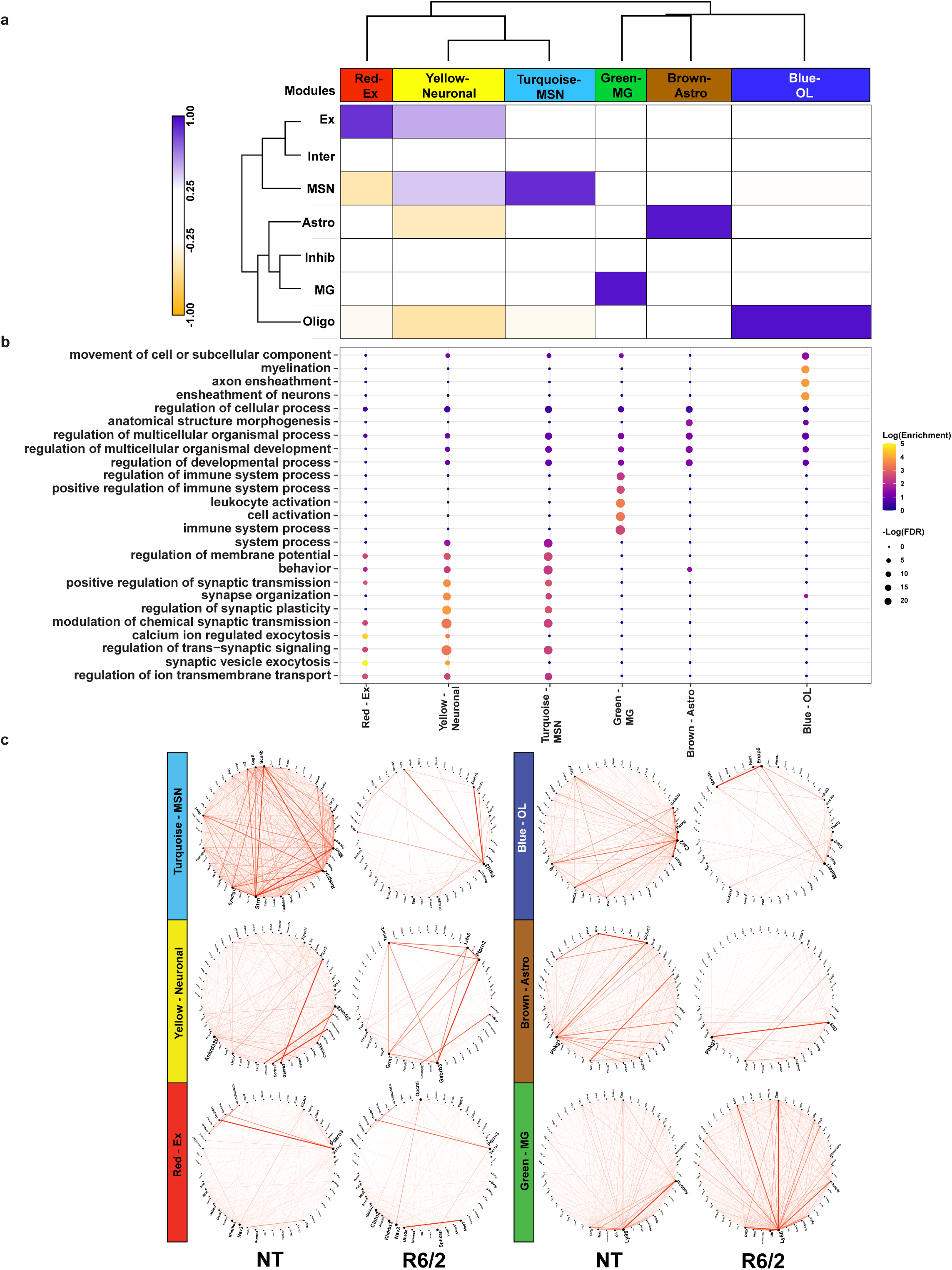
WGCNA analysis of R6/2 mouse snRNAseq data shows cell type-specific changes in network structure. **a**) Dendrogram and correlation heatmap showing cell type-specific co-expression modules. Heatmap shows modules highly correlated with each cell type, dendrogram shows clustering of neuronal module together and glial together. Cell color represents column min (orange) and max (blue). **b**) Top five GO terms per module, showing cell type-specific functional relevance. **c**) Circos plots of the top 50 genes with highest kME in NT mice (**left**) and R6/2 (**right**). Red lines show connectivity between the top 50 genes. Structural differences can be seen between NT and R6/2.

To understand the potential causal connections between these genes and HD, we applied a Bayesian approach to causal network modeling (See Methods) with the combined cell type-specific WGCNA module genes and cell type-specific DEGs as input (**Fig. 4a** **and** **b, Supplementary Fig. 5b-d, Supplementary Table 4**). We explored the MSN and OPC/OL bayes nets (bnets) in detail for two reasons: 1) since MSN are the most studied cell type in HD the bnet should recapitulate previous findings and also reveal both known and novel interactions between known dysregulated genes, providing validation for our approach, and 2) both cell types were the most impacted in our mouse model (total number of DEG) with the OPCs and OLs showing the largest number of DEGs that suggest developmental deficits. The merged NT and R6/2 bnets are shown in **Figs 4a** **and** **b**. We highlight genes representing key drivers (hub genes with high outward centrality, or genes connecting 2 hubs) which are potentially causal regulators of downstream nodes.

**Fig. 4.**
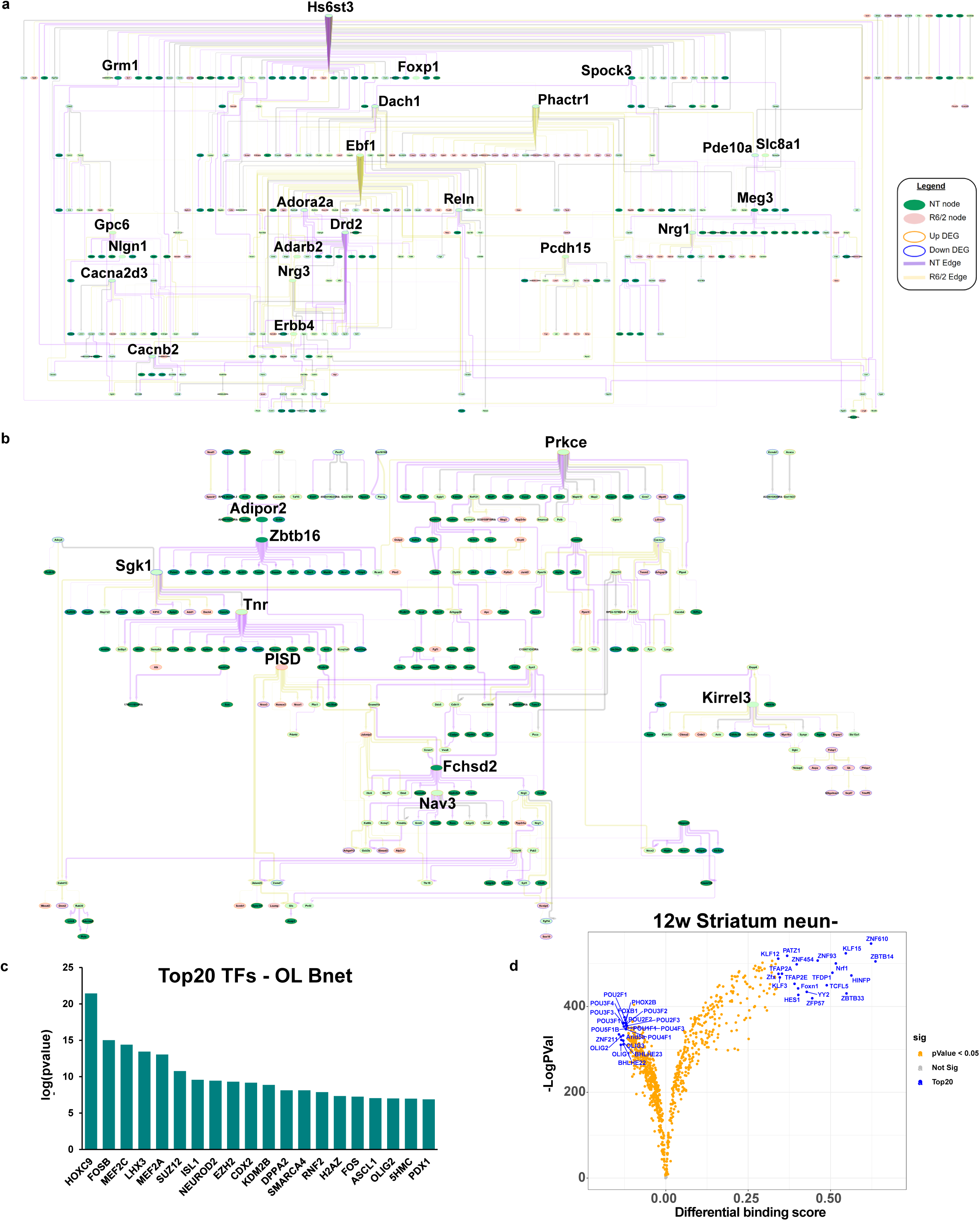
Causal network analysis and ATACseq of glia reveals Prkce, Olig1/2, Sox9/10, and glucose and lipid metabolism as important regulators. **a**) MSN bnet. **b**) OL bnet. **a & b**) Both causal networks are merged from NT and R6/2. If a node and edge existed in both the NT and R6/2 bnets, the NT data (edge weight) were used for plotting. Each bnets shows nodes that exist only in NT or R6/2 and nodes that exist in both, as well as novel edges and edges retained in the R6/2 data. Each bnet was also plotted using a hierarchical structure to show the direction of causal flow. In each plot, genes with a high degree of outward centrality ( >10 outward edges) are highlighted by increased gene name size, as well as genes that connect between two genes that have a high degree of outward centrality. We consider these highlighted genes key drivers of the network. Color scheme is as follows: Edge (purple = NT, yellow = R6/2, grey = both), node fill color (green = NT node, pink = R6/2 node, light green = both), node outline color (orange = upregulated, blue = downregulated). MG, Astro, and Ex neuron bnets are in **Supplementary Fig. 5b-d. c**) LISA analysis of OL causal network gene members, showing the top 20 regulatory transcription factors. **d**) Volcano plot showing differential binding scores, and - log(pvalue) differences of TF binding in open chromatin in 12w NeuN-striatal cells. blue = top20 by differential binding score, orange = pvalue <0.05. 8wStr, cortical, and all NeuN+ data can be found in **Supplementary Fig. 6b**.

#### MSN Network

The MSN bnet includes genes involved in MSN development/identity, function, and genes implicated in HD, including *Ebf1*, a key driver that is lost in the R6/2 bnet (yellow edges) and is involved in striato-nigral MSN development and other genes that interact in both the direct and indirect pathways ^43,44^. Genes of the indirect pathway in D2 MSNs, including *Adora2a, Drd2, and Penk*, were all downregulated and only show NT causal interactions (purple edges), indicating a loss of function of these genes, thus validating the approach ^45^. Furthermore, *Drd2* is a parent node of *Penk*, which is not only a downstream target of Drd2 signaling and dysregulated in HD ^46^, but is transcriptionally regulated by Drd2 expression through dopamine-induced activation ^47^.

#### OPC/OL Network

Based on the extensive dysregulation of OPC and OLs, we next explored the corresponding bnet (**Fig. 4b**) and found *Prkce, Sgk1, Zbtb16* and *Tnr* as key drivers. Prkce is regulated by DAG and Zbtb16 ^48^, a zinc finger binding protein that is involved on OL maturation and myelination, is found downstream of Adipor2, an adiponectin receptor that regulates glucose and lipid metabolism. Downstream of *Zbtb16* is serum- and glucocorticoid-inducible kinase 1 (*Sgk1),* which is normally upregulated in OLs during cellular stress and regulates many ion channels and solute carrier proteins involved in metabolic pathways and glucose uptake (e.g.^49^), such as GLUT1, GLUT4, and glutamate transporters. *Sgk1* is downregulated in R6/2 mice indicating a potential loss of function in HD – see supplementary results for additional validation studies. Exploration of downstream nodes reveals a connection between *Smarca2*, which is a protein in the SWI/SNF family involved in gene expression and chromatin remodeling in OLs, and *Prkce*. Smarca2 (BRM) and Smarca4 (BRG1) play roles in OPC and OL development, including promoting OPC differentiation ^50,51^. The majority of the outward edges from key drivers are NT specific, indicating a loss of causal connection to downstream nodes in the R6/2 mice. Transcription regulator analysis using LISA ^52^ revealed the network is enriched for targets of Smarca4, and Olig2, as well as other regulators previously highlighted for HD, including Suz12, Jun, Fos, and Mefc2 (**Fig. 4c**). These findings suggest an interconnected role of OPC/OL development with lipid and glucose metabolism through *Prkce* and DAG, protein glycosylation, *Adipor2*, and *Sgk1*.

MG, Astro, and Ex neuron bnets are described in the supplementary results.

### ATACseq of glial-enriched nuclei identifies regulators underlying transcriptional pathology in HD glia

To understand the drivers of gene expression changes in non-neuronal cells (e.g. glia) versus neurons, and validate the LISA analysis, we performed ATACseq on NeuN+ and NeuN-sorted nuclei from both the striatum and cortex of the same R6/2 mouse cohort (**Supplementary Fig. 6a).** The neuronal nuclear protein NeuN is localized in nuclei and perinuclear cytoplasm of most of the neurons in the central nervous system. We performed foot printing analysis using TOBIAS ^53^ which revealed developmental changes in the glia-enriched NeuN-data (**Fig. 4d** **(12w striatum) and Supplementary Fig. 6b, and Supplementary Table 5**), and enrichment for immediate early genes in the neuron-enriched NeuN+ data. Among the top 20 TFs in the NeuN-data that showed differential binding between R6/2 and NT we found Sox9 and 10 were significantly decreased in the 8wk striatal data, and Olig1 and 2 decreased in the 12wk striatal data. Interestingly, when all the samples were grouped and we compared the top 20 up and down TFs per an age and region, there was some overlapping TFs between the 12w cortical and both striatal samples, but these were in opposite directions such as Hes1 and Zbtb14 (**Supplementary Fig. 6b & c**). The 8w cortical samples had the least similarities compared to all other regions and ages (**Supplementary Fig. 6b & c**) and showed a number of HOX genes within the top 20 TFs with reduced binding. The cortical data showed differential binding of other known HD genes such as Egr1 and Sp1. NeuN+ cells have some similarities with the NeuN-showing differential binding of Zbtb14 and Hes1, although in opposite direction, in several ages and regions, but also showed an enrichment for immediate early genes Jun, Fos, and Mef2c/b/d (**Supplementary Fig. 6b**).

### Single nucleus RNAseq from HD and control cingulate, caudate, and nucleus accumbens identifies several heterogeneous OL lineage cells and altered maturation states

Given the altered gene expression in OL lineage cells in R6/2 mice, we investigated whether mHTT expression also impacted gene expression in OPCs and OLs in human HD post-mortem tissue. snRNAseq was carried out on 66 samples from 29 donors (3 grade I, 4 grade II, 4 grade III, 3 grade IV, 5 juvenile-onset HD, and 10 matched controls - the demographics of whom are outlined in **Supplementary Table 6**). To define the pathology in different brain regions, we microdissected the cingulate cortex, the caudate, and the nucleus accumbens from frozen brain tissue as outlined in **Fig. 1a** and analyzed the samples using snRNAseq. All major lineages were identified in the 290525 nuclei analyzed. Projection of nuclei in tSNE space shows that nuclei of the same lineages largely occupy neighboring space (**Fig. 1e** **and Supplementary Fig. 1d&e**). Nuclei did not show distinct donor or batch related colocalization in the tSNE space after correcting for batch effects (**Supplementary Fig. 7 a-b**). A violin plot of lineage-specific genes delineated all expected lineages (**Supplementary Fig. 1e**). We detected changes in gene expression in all cell types; for this study we focused on cells of the OL lineage.

We focused on OLs and OPCs (**Fig. 5a-b**) and analyzed 80199 OL and 13844 OPC nuclei in isolation of other lineages. Projecting OL and OPC in their own reduced dimension space (PHATE reduction – see Methods) shows a continuous trajectory from OPCs to OLs, and separation between HD and control nuclei (**Fig. 5a, b**). To examine the differentiation states of OL lineage cells, using well-established methods ^54^, we calculated the relative ordering of cells along a pseudotime dimension calculated based on the PHATE reduction and projected the pseudotime values in the reduced dimension space (**Figure 5c**). OPCs were set as root nodes and therefore had low pseudotime values, while OLs had high values. Similar to our mouse data, examination of pseudotime values per anatomic region in control, grades I-III HD, and Juvenile onset HD nuclei show altered maturation states across brain regions and grade in HD. That is, across all brain regions examined, HD nuclei showed a relatively larger proportion of cells with intermediate pseudotime values compared with controls, which is more pronounced with increasing HD grade, particularly in HD grade 3. Conversely, in juvenile onset HD (HDJ), the effect was less appreciable in the cingulate cortex, and more pronounced in the striatum, with the majority of caudate and accumbens OPCs showing intermediate pseudotime values compared with control nuclei (**Fig. 5d**). In contrast, HDJ OLs do not show demonstrable differences compared with control nuclei base on pseudotime analysis. The results suggest that HD maturation pathology is at least partially progressive with HD grade, and that HDJ maturation pathology affects mainly OPCs.

**Fig. 5.**
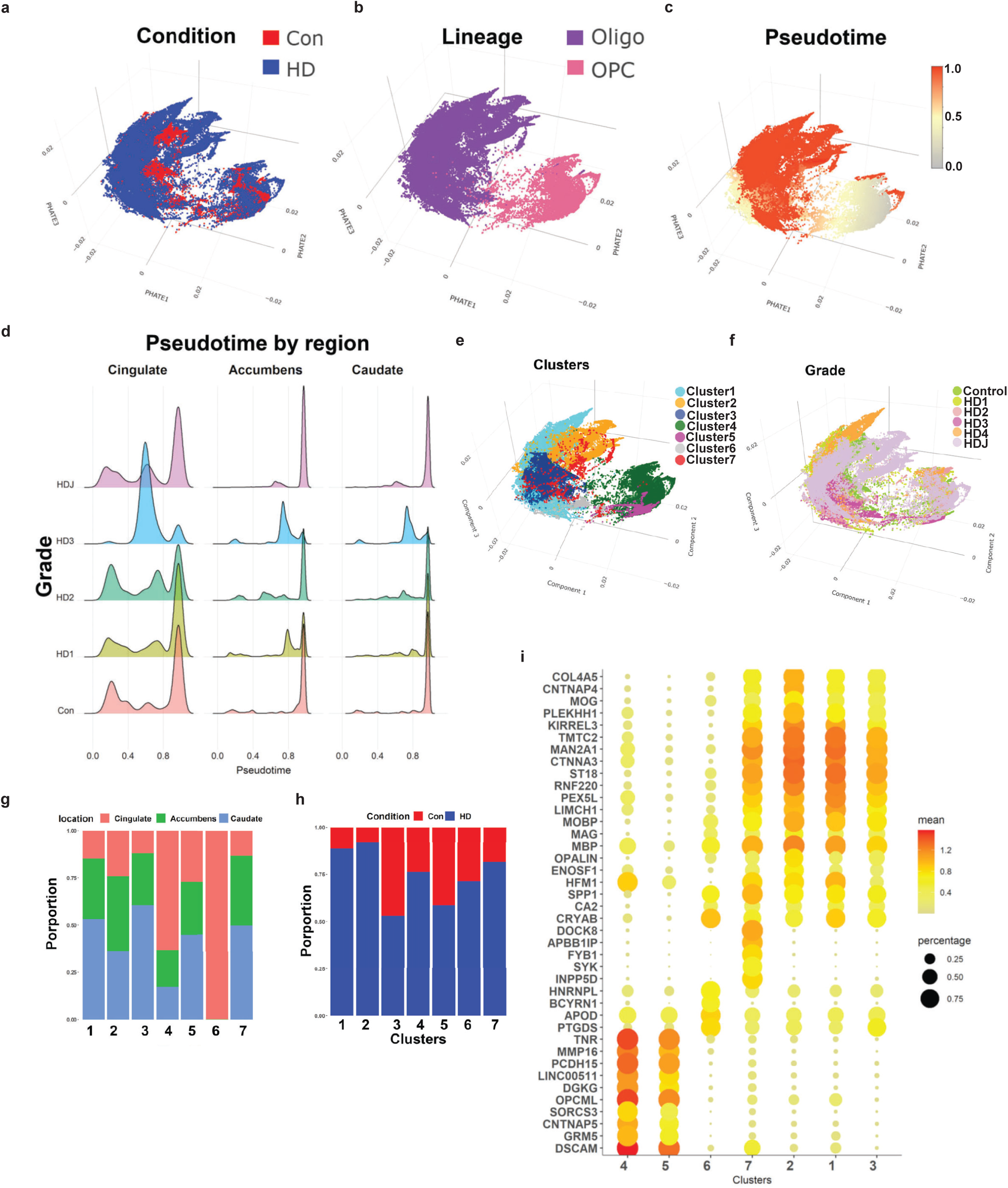
Huntington disease oligodendrocytes are less mature. **a-c, e, f)** Projection of control and HD nuclei in the PHATE dimension color-coded by condition (**a**), lineage (**b**), pseudotime value (**c**), cluster (**e**), and HD grade (**f**). Note that OPCs have the lowest pseudotime values in **c**. **d**) Pseudotime values are shown in histograms across brain region and HD grade. Note that the proportion of nuclei with intermediate pseudotime values is higher in HD, especially grade III. (**g-h**) The relative contribution of anatomic region (**g**) and condition (**h**) to each cluster is shown in bar plots. **i**) Gene expression dot plots showing normalized expression of select cluster marker genes, with color denoting expression levels and circle size denoting the proportion of nuclei expressing the gene of interest.

We next performed unbiased sub-clustering of OL and OPC nuclei using the Levine algorithm and identified 7 sub-clusters (**Fig. 5e**). Most subclusters contained a mix of cells from all three regions (**Fig.5f**) and HD grades (**Fig.5g, h**), although in clusters 4 and 6 most nuclei were derived from the cingulate, and in clusters 1, 3, and 7 caudate nuclei represented the largest proportion (**Fig. 5g**). Most clusters contained mixtures of nuclei from both HD and controls, but a number showed a preponderance of one or the other (**Fig. 5h**) with the caveat that our dataset harbored relatively larger numbers of HD nuclei versus control (Con 17955, HD 76088). With that caveat, Cluster 2 was mostly composed of HDJ nuclei, while cluster 6 was composed of a preponderance of HD3 nuclei (**Supplementary Fig. 7c**). Examination of select gene markers shows that clusters 4 and 5 represent OPCs with relatively high expression of OPC markers *TNR* and *DSCAM* (**Fig. 5i, supplementary Fig. 7d**) and low expression of gene markers for mature OLs. Compared to cluster 5, cluster 4 shows lower expression of OPC genes BCAN, VCAN, PDGFRA, and CSPG4, but a higher proportion of cells with TCF7L2 expression, suggesting this cluster represents differentiation-committed OPCs ^55^ (**Supplementary Fig 7d)**. Conversely, clusters 1, 2, 3, and 7 show relatively high expression of OL genes *CNP, PLP1*, and *MBP* (**Fig. 5i**). Amongst the former, cluster 2 shows the highest expression levels of OPALIN and MOG, suggesting it is most mature (myelinating). Moreover, cluster 7 showed expression of both OL genes (although at comparatively lower levels) and the OPC gene DSCAM and is interpreted as an intermediate state between OL and OPC lineages. Likewise, cluster 6 showed expression of the immature OL gene CA2 as well as other OL genes including APOD, PTGDS, and CRYAB, but not myelin genes. It is thus also interpreted as immature OL. Interestingly, the HD-enriched clusters 1, 2, and 7 showed higher expression levels of KIRREL3 compared with the control-enriched cluster 3. KIRREL3 is a gene shown to be highly expressed in OL residing in chronic inactive lesions of multiple sclerosis^55^. Finally, the HD-caudate predominant myelinating OL Cluster 7 showed relatively high expression of several immune related genes such FYB1, SYK (**Fig. 5i**), APOE, CD74, and C3 (**Supplementary Fig. 7d, Supplementary Table 7**), reminiscent of the immune oligodendroglia described in multiple sclerosis^55^. The cluster markers are provided in **Supplementary Table 7.**

### Differential gene expression analysis reveals further differences between HD and control OLs

We next identified significant DEGs between HD and control OL and OPC nuclei in different regions; the number of significant DEGs unique to and shared by respective anatomic regions is shown in Venn diagrams for OLs (**Fig. 6a, Supplementary Table 8**) and OPCs (**Fig. 6b**, **Supplementary Table 8**). Given that the neurodegeneration is detected in the caudate nucleus at the earliest stages of HD and that pathology in the nucleus accumbens and cortex is typically seen in more advanced disease, we reasoned that comparing DEGs in these regions is informative in the following ways: 1) DEGs that are shared among the caudate, accumbens, and cingulate likely represent pervasive or core transcriptional pathology in different anatomic regions regardless of disease severity. 2) DEGs shared between the relatively preserved nucleus accumbens and less severely affected cingulate cortex likely represent early pathologic alterations that may be compensatory in early stages of the disease and are lost in more advanced stages. This does not preclude the possibility that any number of these DEGs may represent cell-autonomous changes due to mHTT in OL and OPCs. With this insight, examination of significant DEGs in these regions highlights a number of themes; first, myelin related and OL identity genes including MAG, MBP, MOBP, MOG, OPALIN, PLP1, CNP, and OLIG1 and 2 were significantly downregulated in OLs of all areas in HD (**Supplementary Table 8**). This was reflected in a negative enrichment of the GO myelination in HD OL’s across all three brain regions (**Fig. 6c**). Second, multiple heat shock response genes including HSPA1A, HSPH1, HSPA4L, HSP90AA1, HSPB1, HSPA4, and HSPD1 were increased across all anatomic regions, suggesting widespread, pervasive pathology in HD OLs (**Supplementary Table 8**). Multiple metallothionein genes including MT2A, MT3, MT1X, MT1M, and MT1E, as well as heat shock protein encoding genes HSPA1A, HSPA1B, and HSPB1 were increased in all brain regions in HD (**Supplementary Table 8**). SPP1, which is a secreted protein that is increased in demyelination and remyelination ^56^, was also increased in all these regions. CA2, a gene encoding a carbonic anhydrase enzyme expressed in immature OL and mature OLs but not OPCs^57^, was increased in cingulate OLs (validated in **Supplementary Fig. 8b-e**). To determine whether similar metabolic genes were dysregulated in our human OPC and OLs that were found in our mouse data, we overlapped human OPC and OL DEGs with the dysregulated metabolic genes in the 12w striatum data and found a large overlap of with these DEGs (**Supplementary Fig. 8a**) including DGKx, GALNTx genes, PTGDS, and TPK1. In the accumbens and cingulate (**Fig. 6c**), gene ontologies related to nuclear export, RNA binding, RNA splicing, peptidyl-lysine modification, and H3 deacetylation were more significantly enriched. Several DEGs shared between the accumbens and cingulate OLs were related to metabolism, including adipogenesis (ARL4A, COQ3, CHUK, ABCA1, GBE1, and ME1 – increased in HD OLs), fatty acid metabolism (EVOVL2 and PLA2G6 – decreased in HD OLs), and pyruvate metabolism (pyruvate kinase M1/M2 PKM - decreased in HD OLs). These results implicate metabolic pathways, including lipid and glucose metabolism in HD pathology at early stages of neurodegeneration (**Fig. 6c** **and Supplementary Table 8**). The involvement of immune genes we observed in HD-enriched clusters is reflected in the enrichment of immune-related ontologies in the HD OLs DEGs, including NFKB activation and inflammasome (**Fig. 6c** and **Supplementary Table 8**). Analysis of enriched GO in HD OPCs reveals a downregulation of genes related to N-acetyl-galactoseaminyltransferase activity, and an upregulation of stress-related ontologies across the three regions. Similar to the mouse data, we also see terms related to nervous system development, ion channels, and cell adhesion (**Figs. 2a** **and Supplementary Table 8**).

**Fig. 6.**
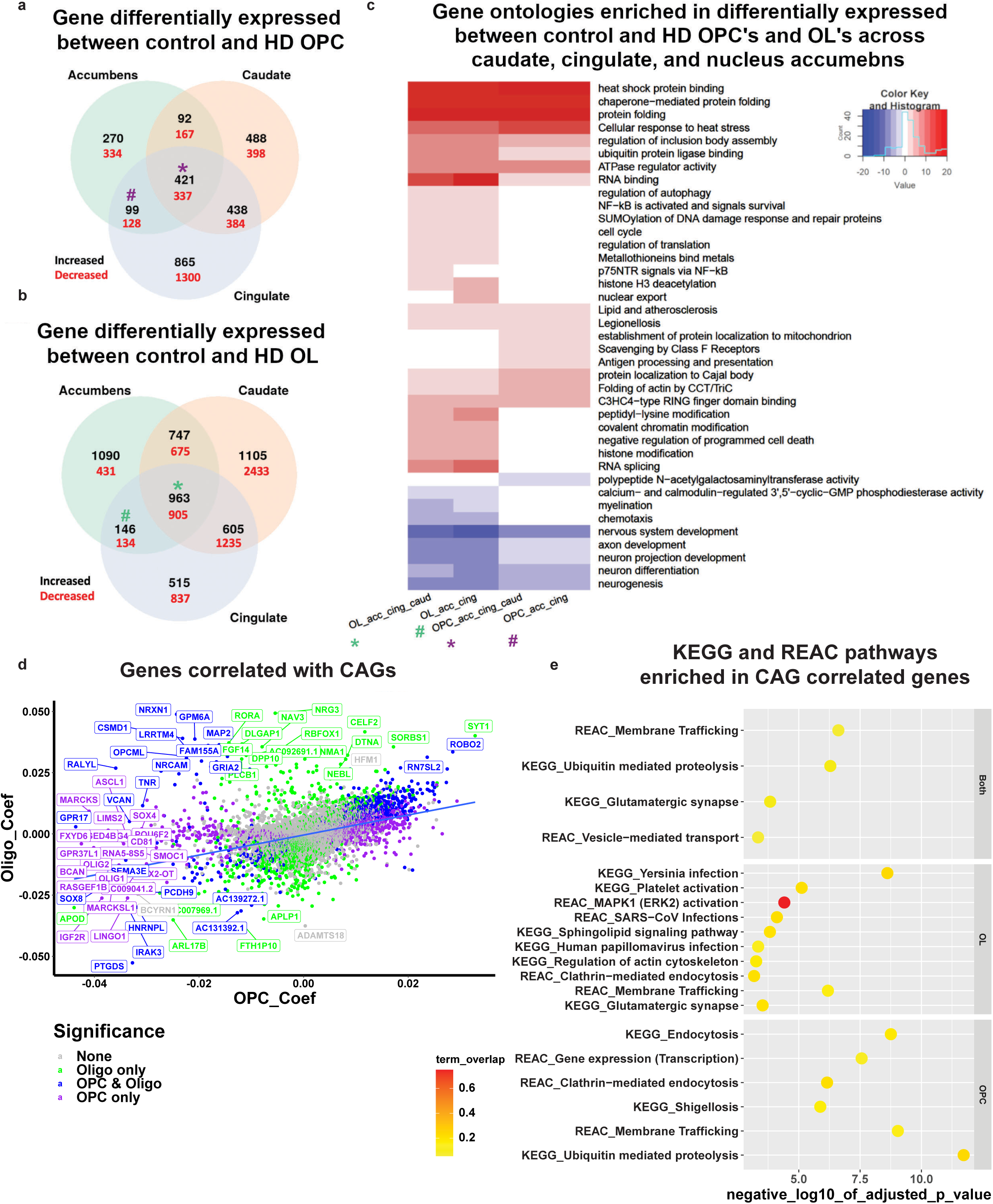
Differential gene expression analysis of HD and control OPCs and OLs. Venn diagram analysis of the DEGs in OPCs (**a**) and oligodendrocytes (**b**). The number of DEGs that are increased (black) or decreased (red) in HD nuclei is highlighted per overlap sector. The stars indicate the DEGs that are shared across all regions, and the # indicates the DEGs shared between the Cingulate and Accumbens. **c**) Gene ontology (GO) term analysis of differentially expressed genes in select sectors of the venn diagrams HD versus control OLs and OPC (from panels a, c). The * and # signs correspond to the DEGs shared across all regions and DEGs shared between accumbens and cingulate OL and OPCs, respectively (purple = OPC DEGs, and green = OL DEGs). The sign of the negative log10 of the adjusted p value indicates the direction of changes; positive sign corresponds to genes increased in HD, and negative sign corresponds to genes decreased in HD. **d**) Scatter plot of the correlation coefficients of genes that correlate with CAG repeats in OPCs (y-axis) and OLs (x-axis). The color of thew genes correspond to whether the coefficient was significant in OLs only (green), OPCs and OLs (blue), or OPCs only (purple). **e**) KEGG and Reactome pathway enrichment analysis of the genes that significantly correlate with CAG repeats in OPCs and OLs (top panel), OLs (middle panel), or OPCs (lower panel). The negative log10 of the adjusted p value is indicated on the x-axis, and the pathways on the y-axis. The color of each circle corresponds to the percentage of overlap between the CAG-correlated genes and the genes in each pathway.

### Dysregulated gene expression is related to numbers of CAG repeats

The length of CAG repeats varied among our donors, and even between regions in the same donor (**Supplementary Table 6**). To determine if any of the OL or OPC genes varied as a function of the numbers of CAG repeats, we conducted a regression analysis with gene expression as response variable and CAG repeats as explanatory variable. We collapsed cells from each sample and used the pseudobulk samples as input for the regression analysis, corrected for batch and brain region and only extracted the significant CAG coefficients (**Supplementary Table 7**). A number of genes showed significant correlations between expression and CAG repeat lengths, some in OPCs or OLs or both (**Fig. 6d**). The graph plots the regression coefficients of each gene in OLs versus OPCs; the upper right quadrant represents genes with positive correlations in both OPCs and OL, the lower left quadrant genes that have negative correlations in both. Among genes with negative correlations in OPCs are transcription factors *OLIG1* and *OLIG2*, *ASCL1*, *SOX2* and *SOX4*, which play roles in OL-lineage development, along with *IGF2R*, suggesting that progression through the OL lineage is further inhibited with longer repeat length. Indeed, OPC lineage genes including *OPCML* and *CSPG4* were negatively correlated with CAG repeat length (**Fig. 6d**). Moreover, *PTGDS*, a cluster 6 marker, had the most negative coefficients in both OPCs and OLs as a function of CAG repeat length, implicating prostaglandin synthesis in the severity of HD pathology. Some of these genes also were identified in our OL bnet as key drivers, including: SGK1, TNR, and NAV3 (**Fig. 4b**). We also investigated KEGG and REAC pathways that were enriched in genes correlated with CAG repeat lengths (**Fig. 6e** and **Supplementary Table 7**). Among the pathways that are enriched in OLs with increasing repeat lengths are those of inflammation, which is more pronounced in human brain, sphingolipid signaling, and ERK2 activation – which is known to control myelination^58^. Both OLs and OPCs show enrichment in genes related to glutamatergic synapses and ubiquitin-mediated proteolysis. When we examined the OL genes with negative coefficients, we found that a number of them are involved in cholesterol metabolism including (DHCR7, DHCR24, ABCA2, and ACAT2 – **Supplementary Table 7**), which further implicates lipid metabolism as central to OL pathology in HD.

### Validation of OL pathology in human HD and mouse data

Many genes that regulate OL maturation or were identified as key regulators were similarly dysregulated in HD patient and mouse data including: MOBP, MAL, CLDN11, MBP, OLIG1, OPALIN, PRKCE, and SMARCA2 (**Fig. 7a**). To confirm dysregulation of key genes PRKCE and TPK1, performed WB analysis. Additional investigation and validation of OL genes and other metabolic genes was also conducted and can be found in the supplemental data and text. Protein levels of PRKCE, and phospho-PRKCE were significantly decreased in the cingulate and caudate of HD brains and the ctx and str in the R6/2 mice (**Fig. 7b-e**). Both species showed an increase in *PRKCE* RNA levels, opposite of the protein data. The ratio of p-PRKCE to PRKCE was not altered though, suggesting that reduction in active PRKCE is related to reduced protein levels (**Fig. 7b-e**).

**Fig. 7.**
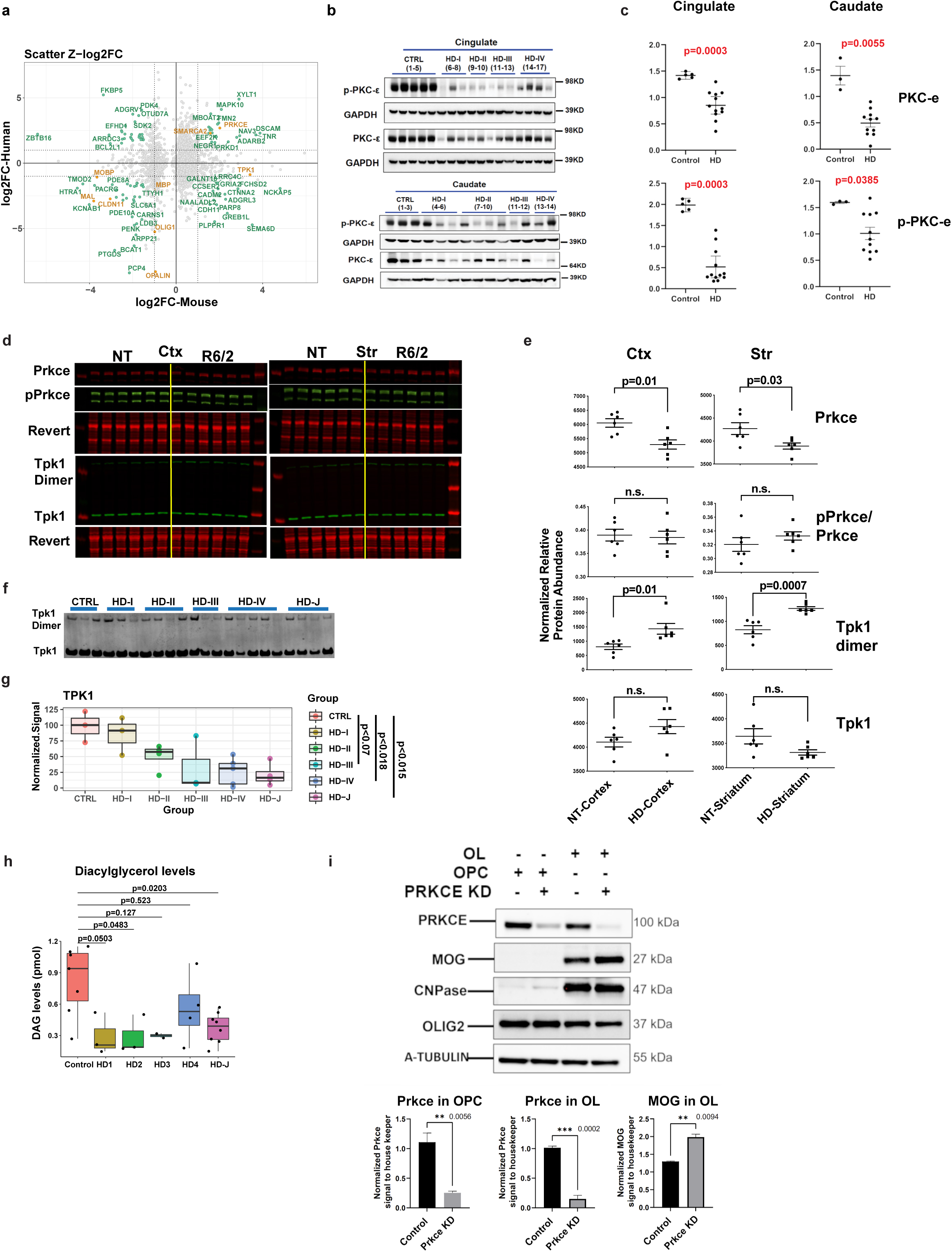
Western, lipidomics, and cellular analyses validates HD differences in TPK1 and PRKCE. **a)** Scatterplots of Z-score log2 fold change values comparing mouse and human data in 12w striatum versus human caudate OL DEGs. Genes with |Z-log2FC| values > 1.5 are highlighted in seafoam green and OL maturation genes are highlighted in orange, showing concordance between species for PRKCE and OL maturation genes, and discordance of TPK1 expression. Seafoam green = genes with absolute value(zlog2FC) differences > 1, Orange = key genes highlighted. **b**) Western blot of PRKCE and phospho-PRKCE in HD and control patient cingulate cortex and caudate. **c**) Quantification of western blow results. Mann Whitney test used for each statistical analysis. Exact p-values: Cingulate: PKCE-0.0003, p-PKCE-0.0003; Caudate: PKCE-0.0055, p-PKCE-0.0385. **d**) Licor images of Prkce, pPrkce, TPK1, and respective revert in R6/2 and NT striatum and cortex. **e**) Quantification of licor results. **f**) Western blot of TPK1 in human caudate samples from juvenile HD, HD grades 1-4, and control patients. **g**) Quantification of human TPK1 data. Statistical analysis was done using a one-way ANONA and Tukey HSD posthoc, comparing control to each adult HD grades (adjusted p=0.979, 0.221, 0.070, 0.018) and control to juvenile HD (p=0.015). **h**) DAG levels quantified from HD and control patient brains showing significant decreased DAG levels in HD brains. **i)** Western blot of PRKCE, MOG, CNPase, OLIG2, and A-Tubulin in OPC and OLs +/- K/D of PRKCE.

Since TPK1 was found to be dysregulated in both mouse (up) and human (down) data at the RNA level in OLs and OPCs, we assessed the protein levels of the monomer and active dimer form of TPK1. **Fig. 7f-g** shows a decrease of TPK1 (monomer and dimer) in HD patient tissue with HD grade 3 & 4 (At adjusted p-value <0.1 for 3, and <0.05 for 4), and in juvenile HD (adjusted p <0.05), consistent with RNA expression data, whereas TPK1 dimer is increased in the R6/2 striatum (**Fig. 7d-e**). The mouse and human data are discordant from each other which may indicate a loss of function of expression in humans and compensatory increase in the mice or other unknown mechanism. Nonetheless, the data confirms that TPK1 is dysregulated in both human HD and murine model of HD.

Given the potential contribution of DAG to OL development and as a substrate of PRKCE – a central hub of the OL causal network, we evaluated DAG levels using lipidomic profiling of control brain versus HD in the cingulate. A significant decrease in DAG levels was observed in juvenile HD brain as well as grade 2 HD brains relative to controls (**Fig. 7h**). These data support the hypothesis that glucose and lipid metabolism, and specifically DAG signaling, potentially through PRKCE, could be playing an important role in the OPC/OL maturation changes we see between HD and control patients. This is further supported by the reduction in TPK1 in HD brains due to the involvement of thiamine in the production of acetyl-CoA, which is then used during DAG formation. Given this finding along with the results demonstrating the reduction of PRKCE in human tissue, together with the causal network analysis placing PRKCE at the top of the OL/OPC network upstream to several maturation genes, we hypothesized that it played an important role in promoting OL differentiation. To test this hypothesis, we knocked down Prkce from primary murine OPC cultures, and differentiated these cells into OLs. The cultures expressed OLIG2, and OLs expressed CNPase. Compared with scrambled siRNA, siRNA specific to Prkce effectively knocked down the protein (**Fig 7i**). Interestingly, the levels of MOG were significantly increased by Prkce knockdown, supporting that the downregulation of Prkce leads to increased OL differentiation. Indicating that loss of PRKCE - as seen in our western blot data - in both human and mouse HD OPCs/OLs would lead to increased OPC commitment to differentiation, an increase in COP cells which we see in our snRNAseq data.

### High Dose thiamine and biotin rescues transcriptional dysregulation in neurons and altered OL and OPC developmental genes in a mouse model of HD

Given that both mouse and human data showed alterations in TPK1 and SLC19A2, and these may regulate PRKCE thorough DAG, we tested whether high doses of thiamine and biotin (T&B) treatment, similar to that used to treat HD-like phenocopy disease such as biotin-responsive basal ganglia disease ^32^, would rescue our observed broad and/or cell type-specific gene expression changes including OL maturation genes. Furthermore, due to the discordant RNA expression changes in our mouse and human data we speculate that the increase in TPK1 was compensatory in the HD mouse model. Considering that TPK1 was only increased at 12w and not 8w, we suspect that these compensatory changes are responding to earlier metabolic changes and tested whether targeting thiamine metabolism at a relatively early timepoint prior to any documented changes in TPK1 expression ^59^, would rescue the dys-maturation. For this study, R6/1 mice were used given symptoms are delayed by several weeks relative to R6/2 mice ^30^, thus allowing a greater window to observe effects of a given treatment. R6/1 and NT mice (8w-old) were treated with vehicle or T&B for 7wks before striatal tissue was collected and analyzed using snRNAseq (**Fig 8a**). MSNs, inhibitory neurons, OPCs, OL, and Astros showed the most DEGs between R6/1 and NT vehicle treated mice (**Supplementary Table 10**). Comparing R6/2 and R6/1 DEGs for each cell type, we found high correlation between HD models and a significant overlap in DEGs, including between OPC and OL maturation genes (**Fig. 8b**) supporting the use of R6/1 mice for the supplementation study. When we evaluated DEGs between R6/1 T&B treated and vehicle treated mice (treatment effect), for each cell type, there was a significant overlap of genes impacted by T&B treatment and genotype DEGs (**Fig. 8b**). **Figure 8c** shows a scatterplot of the overlapping DEG between the T&B treatment effect (R6/1 + T&B vs R6/1 + vehicle) and the genotype DEGs (R6/1 vs NT) for each cell type, which shows significant discordance between the genotype DEGs and the treatment DEGs, indicating rescue of these transcriptional alterations. This translated into a decrease in the number of significant DEGs detected for each cell type ((R6/1 + T&B vs NT) compared to (R6/1 + Vehicle vs NT)), except for the Ex neurons which actually had an increase in DEGs (**Fig. 8d**). Interestingly, the cell types with the most genes rescued by T&B treatment (discordant values) were OL-lineage cells and Adarb2+ interneurons that represent inhibitory neuron subcluster 1 (Inhib1 (**Fig. 8a**)). Based on the reduction of DEGs detected OL, MSNs, Interneurons, Astros, and OPC all had a large decrease in the number of DEGs detected by 115, 176, 378, 129, and 82 DEGs, respectively. Within the OPCs and OLs there was significant rescue of maturation related DEGs *Clnd11* and *Mal*, and a further increase of *Neat1,* which was increased in caudate-parenchymal human HD OLs, and is upregulated during OL maturation. Several genes that correlated with CAG repeat length, e.g. *Ptgds, Phgdh*, and *Tmtc2,* were rescued by T&B treatment. GO enrichment analysis also revealed the molecular functions of the genotype DEGs that were rescued from T&B treatment (**Fig. 8e**). In Astrocytes there was a significant rescue of iron metabolism related genes, Ex neurons showed rescue of neuroligin binding and calcium signaling, and the MSNs showed rescue of cyclic nucleotide phosphodiesterase activity, GABA receptor activity, calcium transport, creatine kinase activity, and electron transport chain genes. Similar to MSNs, the inhibitory neurons showed rescue of calcium related genes, cyclic phosphodiesterase activity, and creatine kinase activity, but also showed unique terms such as glutamate receptor activity, LDL binding, neurotrophic TRK receptor, and fructose binding. Lastly, the OPCs and OLs showed rescue of glutamate receptor activity, RNA binding, creatine kinase, activity, calcium related genes, and GTP binding. These results a) support the hypothesis that metabolic changes in HD contributes to driving cell type-specific transcriptional changes and b) specifically thiamine metabolism deficits may be contributing to OL maturation deficits.

**Fig. 8.**
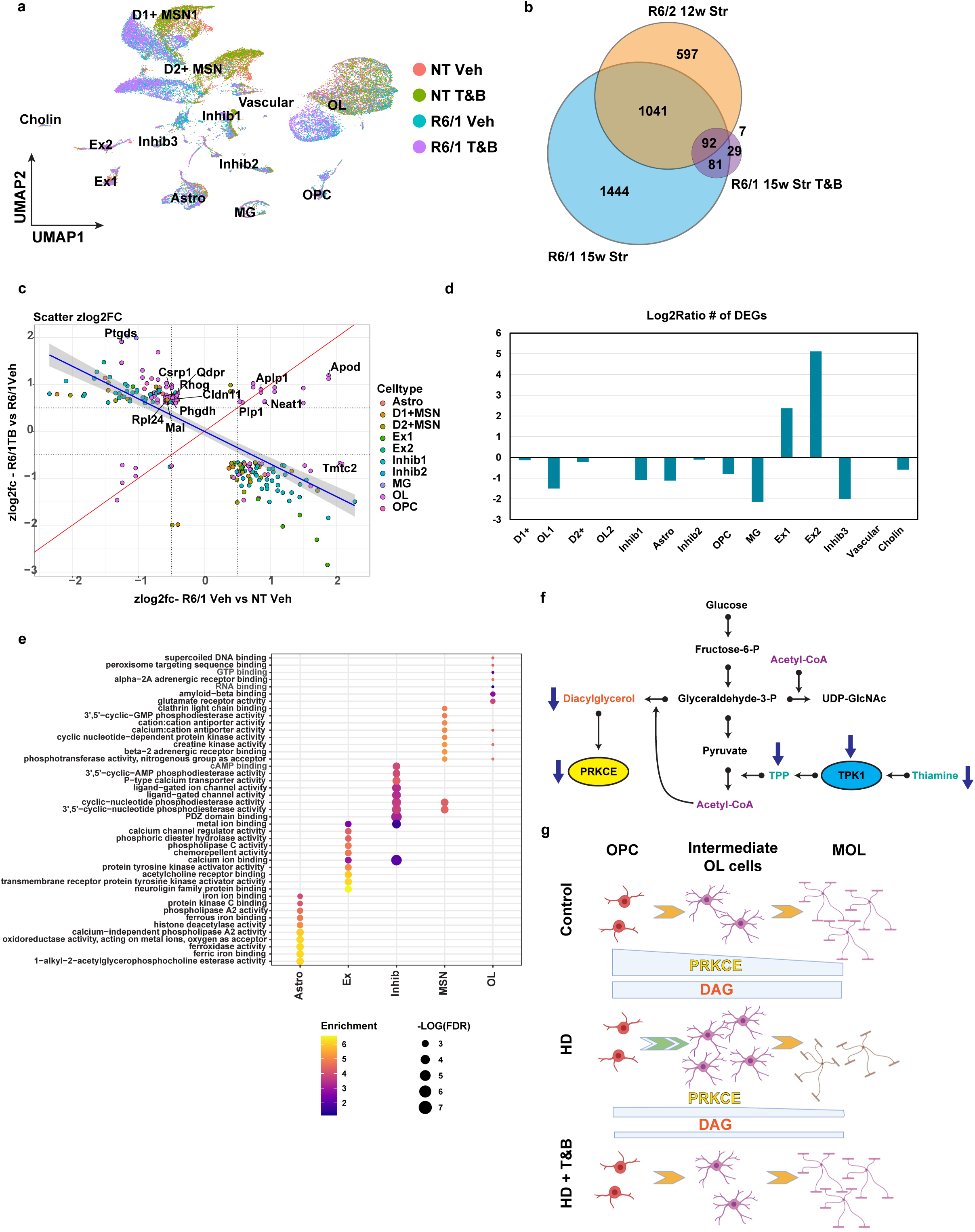
Thiamine and biotin study in R6/1 mice shows rescue of OL maturation DEGs and other cell type DEGs. **a)** UMAP showing the R6/1 and NT mouse data colored by genotype and treatment. **b**) Venn diagram comparing genotype DEGs in 15w R6/1 mice and 12wStr of R6/2 mice against each other and treatment effect DEGs from R6/1 T&B treated versus vehicle. **c**) Scatterplot showing Z-score log2FC of all genes overlapping between genotype and treatment effect DEGs. Colored by cell type origin. OL and Inhib1 neurons show the most rescued DEGs. Quadrants 1 and 3 represent rescue of expression and 2 and 4 represent exacerbation. **d**) Barplot showing the log2ratio of the number of significant DEGs comparing R61 vehicle versus NT vehicle to R6/1 T&B versus NT vehicle. **e**) Top 10 GO terms of overlapping DEGs per cell type (R61 vehicle versus NT vehicle to R6/1 T&B versus NT vehicle). **f**) Illustration of metabolic pathways impacted in HD. **g**) Illustration showing how PRKCE and DAG levels regulate OPC commitment to differentiation and MOL maturation in control and HD, and how T&B treatment rescues maturation impairments.

## Discussion

The studies above describe a systematic and in-depth analysis of single cell transcriptomics of HD mouse models and human patient brains leveraging causal network modeling (CNM) to implicate key drivers of gene expression pathology. Using snRNAseq, we identified dysregulated genes across multiple cell types and cell type-specific changes that may drive the functional changes seen in each cell type. In addition to specific changes in neurons, specifically D1 and D2 MSNs, a large number of gene expression changes in the OL lineage related to development and maturation processes were identified. We defined a progressive dys-maturation phenotype that spans multiple brain regions in both human and mouse HD. CNM identified potential key genes and molecules with putative causal roles in cell type-specific alterations, several of which were connected to metabolic functions, cell maturation, and OL/OPC-identity genes. This includes PRKCE that causally interacts with many other genes in our OPC/OL bnet, including SMARCA2 and OLIG2 targets important in OL maturation. Functional studies validated PRKCE’s role in promoting OL maturation. Our ATACseq data provided further validation demonstrating decreased accessibility for genes regulated by known OL developmental TFs (SOX9 and 10, OLIG1 and 2, and ASCL1)^60^, further implicating OL differentiation in HD pathology. These data provided a framework to build targeted therapeutics, as illustrated by treatment with T&B that restored many of the maturation and transcriptional deficits and providing further validation of the approach.

Recent single nuclei studies identified common and cell type-specific transcriptional alterations in R6/2 and Q175 HD mouse models that were recapitulated in postmortem HD human caudate and putamen ^14,61^, showing cell-type specific alterations in HD. In MSNs, mitochondrial dysfunction underlay a detrimental innate immune response ^14^. Striatal OLs showed decreased expression of several markers, however, the correlation between mouse and human OL signatures was low in this case. Here, we show that OLs are increased in the human cingulate and caudate, and mouse and human OL show similar transcriptional dysregulation and reduced maturation. HD oligodendrocytes are transcriptionally immature across multiple human and mouse brain regions. The fact that this phenotype spans the severely affected caudate, moderately involved cingulate, and the relatively preserved nucleus accumbens suggests that the deficits are independent of disease severity or anatomic region. Nonetheless, our data shows that impaired OL maturation is progressive with HD grade, and that in juvenile-onset HD, the maturation deficits largely involve OPCs. This was supported by ATACseq results demonstrating reduced binding of OL developmental TFs.

Previous studies have suggested that the dysmaturity of HD OLs may also represent an inability to respond to the normal turnover of myelin or dedifferentiation. If the accumulation of mHTT downregulates the transcription of myelin genes, it may inhibit the ability of already myelinating OLs to produce myelin components during their normal turnover. Huang et al. showed that mHTT binds to MYRF and downregulates myelin genes ^17^. MYRF is positively regulated by CHD7, which is regulated by OLIG2^50^– a master regulator of OL identity and a gene our results implicate in HD pathology. While MYRF appears to play a role in the abnormal function of mature OLs, we also suggest that OL defects start earlier during OL development and maturation from OPCs, which is consistent with previous studies^23^. In our human data, this finding was most pronounced in juvenile-onset HD, where maturation deficits appear to almost entirely involve OPCs and not OLs - based on pseudotime analysis. Our results support a model where OPC commitment to differentiation is increased in HD, a process facilitated by downregulation of PRKCE. OL maturation is hampered in HD, as demonstrated in the literature, through mechanisms possibly involving dysfunction of MYRF^17^. The difference between juvenile-onset and adult-onset HD is intriguing. We speculate this may arise from the larger CAG repeat lengths in juvenile-onset HD, and the fact that HTT is expressed more highly in OPCs compared to OLs ^62^. That said, we further describe a progressive pathology in OL differentiation that appears more pronounced with HD grade. Thus, OL-lineage pathology in HD is likely both developmental and progressive.

Metabolic disturbances in HD are hypothesized to directly lead to cellular distress, but less is known about their contributions towards epigenetic regulation, transcriptional deficits, and impact on cell maturation and identity. Both mouse and human snRNAseq data show dysregulation of key genes related to glucose and lipid metabolism that include genes that are within or downstream of several key metabolic pathways, including glycolysis, DAG, the hexosamine and protein glycosylation pathways. A recent study demonstrated that accumulation of unsaturated sterols in OPCs drives their differentiation into OLs, implicating lipid metabolism as functioning in OL differentiation, and not only as generating myelin building blocks^63^. Cholesterol metabolism was implicated in HD pathology by several groups^64–71^. Additionally, DAG lipids which activate PRKCE were decreased in HD brains. Interestingly, protein kinase C signaling has been shown to be important to OPC differentiation, and myelination ^72–75^. We found PRKCE levels to be decreased in HD, and that downregulating PRKCE in OPCs in vitro leads to increased differentiation of OLs. Further determination of the mechanism underlying these findings is the subject of future studies. Moreover, appropriate glucose metabolism is critical for the proper development and function of OLs, as OPCs transition to myelinating OLs ^76–79^. Finally, thiamine metabolism is linked to oligodendrocyte differentiation based on evidence from deficient pyruvate dehydrogenase function in humans, which is known to cause structural white matter abnormalities ^80^, and experimental evidence from pyruvate-dehydrogenase deficient mice, which show a reduction of O4-positive OL/OPCs ^81^.

A highly dysregulated gene and the most common DEG in the R6/2 12w striatal data, TPK1 regulates conversion of thiamine to thiamine-pyrophosphate (TPP), a cofactor required for the conversion of pyruvate to acetyl-CoA, by alpha-ketoglutarate dehydrogenase in the TCA cycle and by ketolase in the pentose phosphate pathway, the latter being active in OL cultures and important for myelinating OLs ^82^. Acetyl-CoA links metabolic processes to many epigenetic regulators of transcriptional control as it is used for histone acetylation, in the TCA cycle for energy and feeds metabolites into DNA and histone methylation, and in the generation of both DAG and UDP-GlcNAc, for PRKCE signaling and use by OGT for protein glycosylation (**Fig. 8f**). Interestingly, mutations in TPK1 are linked to Thiamine Metabolism Dysfunction Syndrome 5, which pheno-copies HD, and mutations in thiamine transporters such as SLC19A3 lead to biotin responsive basal ganglia disease ^83^ which is treated with high T&B supplementation. Driven by our findings and similarities to other human disorders, we evaluated T&B treatment as a therapeutic strategy to reverse HD pathology in R6/1 mice. We hypothesized that TPK1 shows a compensatory increase in HD mice at later ages, responding to earlier metabolic changes. We tested this hypothesis by treating relatively pre-symptomatic R6/1 HD mice. Several transcriptional pathologies in HD were rescued by high dose T&B, suggesting promise as a potential treatment strategy. Excitingly, during the course of our study, a separate study was published showing a decrease in SLC19A3 and TPP in HD patients and in both R6/1 and zQ175 mice^84^. High dose T&B treatment produced both increased thiamine levels in the brain and CSF and behavioral rescue in R6/1 mice as early as 13 weeks. Our snRNAseq data revealed that R6/1 mice show maturation and loss of cell identity genes similar to the R6/2 model and that treatment with T&B in the R6/1 mice, prior to TPK1 or SLC19A3 RNA changes, not only rescued a significant portion of dysregulated genes, including neuronal, but also specifically rescued expression of a specific subtype of inhibitory neurons and OPC and OL maturation genes. Furthermore, there was a reduction in the total number of significant DEGs in all cell types, except for in Ex neurons which may be compensatory changes due to the discordant levels in the genotype and treatment effects, but this requires further study outside the scope of this work. These data provide validation of the two studies and additional mechanistic insight that rescue by T&B likely acts in part through rescue of transcriptional deficits in a subpopulation of inhibitory neurons expressing ADARB2, and of OLs. Specifically rescuing many genes involved in HD pathogenesis such as iron metabolism in astrocytes, calcium and phosphodiesterase signaling and activity in neurons, and maturation genes in OLs. Our data suggests that OL maturation impairments may be driven, in part, by thiamine metabolism and changes in the binding of TFs that regulate OL maturation, including Sox9 and 10 and Olig1 and2. Furthermore, HD OPCs seem to have increased commitment into COP and immature OL which could be driven by decreased DAG and PRKCE, which is rescued by T&B treatment (**Fig. 8g**). It also further supports T&B as a viable treatment for HD, now undergoing a clinical trial in Spain (https://clinicaltrials.gov/ct2/show/NCT04478734), and supports the utility of using single cell approaches to guide therapeutic target identification and evaluation.

## Online Methods

### Mice

All experimental procedures were in accordance with the Guide for the Care and Use of Laboratory Animals of the NIH and animal protocols were approved by Institutional Animal Care and Use Committees at the University of California Irvine (UCI), an AAALAC accredited institution. R6/1 and R6/2 mice have been described elsewhere in detail ^30^. For the study using R6/2 mice , 10 five-week-old R6/2 and non-transgenic (NT) male mice were purchased from Jackson Laboratories and aged to 8 or 12 weeks. For the thiamine/biotin study using R6/1 mice, 10 five-week-old R6/1 and NT male and female mice were purchased from Jackson Laboratories. R6/1 mice (5/grp) were given a daily dose of combined 50mg/kg thiamine and 20mg/kg biotin (Caymen, Ann Arbor, MI) or vehicle (PBS) I.P. beginning at age 8 weeks, treated for 7 weeks, then euthanized at age 15 weeks. All mice were housed in groups of up to five animals/cage under a 12-hr light/dark cycle with ad libitum access to chow and water. Mice were euthanized by pentobarbital overdose and perfused with 0.01 M PBS. Striatum and cerebral cortex were dissected out of each hemisphere and flash-frozen for snRNAseq or biochemical analysis.

### Single nuclei RNAseq

#### Mouse

Single nuclei were isolated from ½ hemisphere full striatal or full cortex in Nuclei EZ Lysis buffer (Cat#NUC101-1KT, Sigma-Aldrich) and incubated for 5 min. Samples were passed through a 70μm filter and incubated in additional lysis buffer for 5 min and centrifuged at 500 g for 5 min at 4°C before two washes in Nuclei Wash and Resuspension buffer (1xPBS, 1% BSA, 0.2U/μl RNase inhibitor). Nuclei were FACS sorted using DAPI to further isolate single nuclei and remove additional cellular debris. These nuclei were run on the 10x Chromium Single cell 3’ gene expression v3 platform. Libraries were QCed and sequenced on the NovaSeq 6000 using 30 bases for read 1 and 98 bases for read2, ß to obtain >=50K reads per a cell. A total of 109,053 cells with 6.1 billion reads were sequenced for the 24 samples with on average 4544 cells per sample with ∼55.6K reads each. Alignment was done using the CellRanger pipeline v3.1.0 (10X Genomics https://github.com/10XGenomics/cellranger) to a custom pre-mRNA transcriptome built from refdata-cellranger-mm10-1.2.0 transcriptome using cellRanger mkref. UMI Count matrices were generated from BAM files using default parameters of cellRanger count command. The gene barcode matrices for each sample were imported into R using the Read10X function in the Seurat R package ^85^ (v3.1.5). Replicates were combined using cellRanger aggr.

#### Human

Dissection of the cingulate cortex, caudate nucleus, and nucleus accumbens from frozen Postmortem specimens was performed on material procured and preserved from autopsies on control as well as grade II and grade III HD. These samples were obtained from the New York Brain Bank. All cases had RNA integrity numbers of > 7. Brain tissue measuring ∼ 5 × 4 × 3 mm were dissected on a dry ice cooled stage and processed immediately as described below. A Table of the cases and controls used is provided in **Supplementary Table 4**. Nuclei were isolated as described in. Briefly, brain tissue was homogenized in a Dounce homogenizer with 12–15 strokes of the loose pestle and 12–15 strokes of the tight pestle on ice in a Triton X-100 based, sucrose containing buffer. The suspension from each sample was filtered through a BD Falcon tubes with a cell strainer caps (Becton Dickinson, cat. no. 352235), washed, re-filtered, washed, followed by a cleanup step using iodixanol gradient centrifugation as described in ^76^. The nuclear pellet was then re-suspended in 1% BSA in nuclease-free PBS (containing RNAse inhibitors) and titrated to 600-1200 nuclei/μl. The nuclear suspensions were processed by the Chromium Controller (10x Genomics) using single Cell 3′ Reagent Kit v2 or v3 (Chromium Single Cell 3′ Library & Gel Bead Kit v2/v3, catalog number PN-1000075; Chromium Single Cell A Chip Kit, 48 runs, catalog number: 120236; 10x Genomics). *Sequencing and alignment:* Sequencing of the snRNAseq libraries was done on Illumina NOVAseq 6000 platformV4 150 bp paired end reads. Alignment was done using the CellRanger pipeline (10X Genomics) to GRCh38.p12 (refdata-cellrangerGRCh38–1.2.0 file provided by 10x genomics). Count matrices were generated from BAM files using default parameters of the DropEst pipeline ^86^.

### QC and filtering

#### Mouse

Based on the distribution of number of genes detected in each cell and the distribution of number of UMIs, nuclei with less than 200 genes or more than 6000 genes were excluded from the downstream analyses. Nuclei with percent mitochondrial reads aligning to mitochondria genes of more than 2% were excluded. UMI counts were then normalized in Seurat 3.0 and top 2000 highly variable genes were identified using FindVariableFeatures function with variance stabilization transformation (VST).

#### Human

To remove low quality cells, we first used the combined quality calls from the CellRanger algorithm as well as the DropEst algorithm. This allowed us to retain more high quality nuclei than either algorithm alone. Data QC was done using the scater package ^87^. Nuclei with percent exonic reads from all reads in the range of less 75% were included. Nuclei with percent mitochondrial reads aligning to mitochondria genes of more than 14% were excluded. Genes were filtered by keeping features with > 10 counts per row in at least in 31 cells. A temporary count slot was created by decontaminating the counts from ambient RNA by calling decontX() function with default parameters in R ^88^. These counts were used for downstream clustering, but not differential gene expression analysis.

### Combining multiple datasets from different sequencing batches and count normalization

Using the R package Seurat (version 4.06)^89^, the datasets were merged after controlling for sequencing batches (four batches). We integrated the lognormalized and scaled datasets in Harmony version 0.1. The Harmony reductions were then added to the merged Seurat object containing all datasets. The merged object was normalized using SCTransform function in Seurat accounting for batch and percentage mitochondrial reads ^90^.

### Dimension reduction and clustering

#### Mouse

Based on the elbow plot, top 20 PCs were retained for seurat objects with all cell types and 15 for the OPC and oligo analysis. These PCs were used in the downstream unsupervised clustering using a shared nearest neighbor Louvain modularity optimization to identify clusters of cells of the same type. Some of the identified clusters were comprised of multiple cell types, therefore we subclustered these cells for further downstream DEG generation and analysis (**Supplementary Fig. 1a**).

#### Human

Pre-clustering of nuclei was done in Seurat using the shared nearest neighbor smart local moving algorithm ^91^ after using the iNMF or UMAP reducions, and calling FindClusters(… , algorithm=3,method=“igraph”, n.iter = 100, …). Several resolution and k options were trialed to select the option with the largest number of pre-clusters with the high lineage purity. Lineage identity was determined for each cluster using was done using geneset enrichment analysis of lineage markers ^92^ and by inspecting cluster markers generated by scran::findmarkers(direction=”up”) function ^93^. We also depended on the cell_classifier tool we previously used ^94^. Pre-clusters with mixed identities based on enrichment of multiple lineage genes were sub-clustered iteratively until all pre-clusters showed pure identities which we combine into lineages (Astrocytes, neurons, oligodendrocytes, myeloid, endothelial, OPCs, and ependymal cells). Sub-clustering of select pre-clusters was done as needed to get the lineage-pure small clusters. We next combined the clusters of the same lineage to call the lineages presented in Fig. 1e.

After getting pure OL and OPCs, a new object from these cells only was created in monocle3. Corpus callosum cells were removed, because no HD corpus callosum samples were included in the dataset. Filtering lowly expressed genes yielded 16955 genes. The SCT normalized counts were used to reduce the dimensions using the PHATE function ^95^ in R correcting for batch (using the mutual nearest neighbor option), and using the following parameters: KNN= 5, Dim=3, Decay=50, T=10. Clustering was done in monocle3 utilizing the three PHATE reductions as input using the Levine algorithm.

### Cluster annotation and differential gene expression

#### Mouse

Unsupervised clustering was done using shared nearest neighbor Louvain modularity optimization. For each cluster, we used multiple cell type-specific marker genes that have been previously described in the literature to determine cell type/state identity. Exemplary genes used as markers for major cell types are shown in **Supplementary Fig. 1**. Differentially expressed genes between different clusters, ages or disease groups were identified using Wilcoxon Rank Sum test on genes that are expressed in at least 25% of the group. Further sub-clustering was conducted on some of the main clusters due to mixed cell types represented in that cluster, e.g. oligodendrocyte progenitors (OPCs) and premyelinating oligodendrocytes and astrocytes with vascular cells. Specifically, for subclustered OPCs and OL, OL-lineage, annotations were used from Marques et al. ^37^ by looking at gene expression for marker genes identified in that study. These annotations were then collapsed into OPC and OL groups for ease of reference and consistency with human OPC and OL cells. Cluster and DEG analyses were conducted on each region and age for HD versus NT independently and, combined where noted that the cells were integrated together across region and age.

#### Human

Differentially expressed genes (DEGs) between HD and control per anatomic region in OL and OPC separately were identified using EdgeR glmQLFTest adjusting for sequencing batch and using an FDR cutoff of 25% (9). The raw counts were used here, not the decontaminated counts. Retrieving the top 3000 differentially expressed genes resulted in adjusted p values less than 0.05, which were considered significant and were used for downstream analysis.

The CAG gene correlation analysis was conducted through the R package limma (version 3.14). Samples for the analysis were prepared using a pseudo-bulk approach. Gene expression data for each donor at a specific region were summed up together respectively to create pseudo-samples for the correlation analysis. Each pseudo donor-region sample were then log normalized and scaled using Seurat’s NormalizeData function (version 4.06) for optimal performance in limma. The covariates accounted for in the design matrix between samples included age and gender. Lastly, a row in the design matrix included the CAG repeats for each donor-region sample. The weights of the model were determined using limma’s lmFit with the arguments of the function including the pseudo-bulk donor region expression data and the design matrix as described above.

### Pseudotime trajectory analysis using Monocle3

#### Mouse

For oligodendrocyte developmental trajectory assessment, cells that were identified as OPC and OL lineage were used to create a separate Seurat object using SubsetData function on raw counts. Pseudotime analysis was conducted on the integrated data across all regions and ages.

#### Human

Pseudotime analysis was done using monocle3 employing the three PHATE dimensions to learn the principal graph using the following parameters: use_partition = F, learn_graph_control = list(euclidean_distance_ratio=0.5, geodesic_distance_ratio=0.7, minimal_branch_len=100, orthogonal_proj_tip=TRUE, rann.k=100), close_loop = F). The root nodes were set as OPC cells. Grade 4 cases were excluded because after filtering low quality cells, two samples had very few OPCs after removing low quality cells and doublets.

### ATACseq

#### Isolation of NeuN+ and NeuN- nuclei

The pulverized tissue was resuspended in 2ml NEB buffer (320mM sucrose, 10mM Tris-HCl pH 8, 5mM CaCl2, 3mM MgAc2, 0.1mM EDTA, 0.1% Triton supplemented with protease inhibitors (Roche, 11836170001) and transferred through 40μm tissue strainer, followed by 5min centrifugation at 600xg at 4C. The pellet was resuspended in 1 ml HS buffer (1.8M sucrose, 10mM Tris-HCl pH 8, 1mM MgCl2 and Proteinase inhibitors) and centrifuged for 20 min at 16,000xg at 4C. The nuclei containing pellet was resuspended in blocking buffer (PBS with 0.5% BSA, 5% Normal Goat Serum and Proteinase Inhibitors) and labeled with anti NeuN-PE antibody (1:100 dilution, Millipore, FCMAB317PE) and with Hoechst (1:2000 dilution, Invitrogen, H3570) for 30min. The nuclei were filtered through 40μm mesh and sorted using BD FACSAria™ with gates set to separate NeuN+ and NeuN- single nuclei populations. The nuclei were collected in tubes pre-coated with 1%BSA and sucrose was added to the sorted nuclei to a final concentration of 0.32M followed by 15min incubation on ice to stabilize the nuclei after sorting. The ATAC-seq was performed as described in Corces et al ^96^. Briefly, 50000 sorted nuclei were transferred to tubes and pelleted by centrifugation at 2000Xg for 15 min. The pellet was resuspended in transposition reaction mix (25μl 2× TD buffer, 2μl transposase, 17μl PBS, 0.5μl 1% digitonin, 0.5μl 10% Tween-20, 5μl water) and incubated at 37C for 30min following by clean up with Zymo DNA Clean and Concentrator kit (Zymo D4004). Illumina adapters were added by PCR to generate sequencing libraries as previously described. The ATAC-seq libraries were sequenced on an Illumina HiSeq 2000 for single-end 50-bp reads. Fastq files were aligned to the mm10 genome using Bowtie2 and paramaters previously described in Smith-Gearter *et al.* 2020 ^97^.

#### Footprinting Analysis

We used TOBIAS software (REF: https://doi.org/10.1038/s41467-020-18035-1) for footprinting analysis of ATAC-seq data. Briefly, aligned BAM files were used to call accessible regions (peaks) using MACS2 using the following parameters: --nomodel --shift -100 --extsize 200 --broad. Peaks from all the samples across all conditions were merged to a set of union peaks using bedtools merge. TF motifs were downloaded from JASPAR CORE 2022 database. TOBIAS software robustly performs all steps of footprinting analysis including Tn5 bias correction, footprinting, and comparison between conditions and has been shown to outperform other common methods of footprinting. TOBIAS also calculates TF binding on a global level across all sites as well as the locus-specific level using JASPAR motif data.

### Gene Ontology, KEGG Pathway, and TF enrichment analyses

#### Mouse

DEGs, gene modules members, and bnet gene members were used for further analyses using GOrilla for gene ontology enrichment analyses, KEGG pathway analysis, and LISA for TF enrichment analysis.

#### Human

Gene Ontology term enrichment analysis was done in gProfiler2 package in R ^92^. The results of edgeR DEG was used as input and the following options: (ordered_query = T, significant = T, exclude_iea = T, underrep = F, evcodes = F, region_query = F, max_p_value = 1, min_set_size = 0, max_set_size = 100, min_isect_size = 5, correction_method = “gSCS”). Statistical significance was determined using the more conservative gSCS method 38 yielding adjusted p values. Terms with adjusted p values < 0.05 were considered significant. The terms shown in the Figs. are selected based on ordering the results based on negative_log10_of_adjusted_p_value followed by the ratio of the shared of number of genes enriched in a term to that of the total number of genes in the GO term (desc(intersection_size/term_size)).

### Network modeling

#### Mouse

Weighted gene co-expression network analysis (WGCNA) ^39^)was used to identify gene network modules from the mouse snRNAseq data. Normalized count data from Seurat 3.0 were first used for feature selection, filtering all genes without at least 1 count in 25% of all cells. Co-expression networks were then generated for NT data using WGCNA. Correlative module-trait relationships were used to identify gene network modules that had high correlation with specific cell types used as input, and module preservation statistics were used to assess recapitulation of gene networks in R6/2 data. *Bayesian network modeling.* To identify causal relationships between cell type-specific gene subnetwork we used a bayesian network modeling approach using the R package BNLearn (Scutari, M. (2010). Learning Bayesian Networks with the bnlearn R Package. Journal of Statistical Software, 35(3), 1–22. https://doi.org/10.18637/jss.v035.i03). Probabilistic graphical modeling has been previously used to assess causal relationships between genes/proteins with great success in recapitulating known biological pathway interactions from single cell data ^98^. Our approach took advantage of the co-expressed gene networks we previously identified to try and find causal relationships amongst these genes. To better interpret our data we chose to use input data from individual cell types, which were identified to be most correlated with each individual gene network module. The resulting causal network would be cell type-specific and easier to biological interpret. Features were chosen based on their inclusion within these gene modules and additionally genes were added based on differential expression between R6/2 and NT mice for each cell type-gene network module pair. E.g. We identified that the turquoise gene network module most highly correlated with our MSNs, these genes and DEGs found in both D1 and D2 MSNs were used as input from both 8 and 12w striatal and cortical data. HD and NT networks were separately generated to identify changes in network structure between disease and control. No priors were used as input for the structure learning. Using this input we constructed our Bayesian networks with a bootstrap approach using 50% of samples and 200 rounds. Due to the spasticity of single nuclei data, even after gene filtering, we chose to use an interval method for discretization, factoring input data into 3 breaks. For structure learning we utilized Bayesian Dirichlet likelihood-equivalence scoring and a hill-climbing algorithm for searching for network structures. An average network was generated from each output where the strength and direction (empirical frequency computed from the probability of each edges’ existence and direction) of each causal edge were greater than or equal to 0.85 and 0.5, respectively. HD and control networks were then merged to identify changes in network structure, novel nodes and edges.

### Primary oligodendrocyte culture

Mouse primary oligodendrocyte precursor cells (OPCs) were isolated with immunopanning as described previously ^99^. Briefly, cerebral cortices from C57BL/6 pups at P7 were digested in papain solution for 20min at 37°C, followed by titration and filtration. Cells were then sequentially incubated in three immunopanning dishes (2 negative selections with BSL1, followed by 1 positive selection with anti-mouse CD140a antibody (BD Bioscience, 558774). After positive selection, OPCs were trypsinized, plated onto PDL-coated culture dishes with SATO medium supplemented with growth factors (10 ng/mL PDGF-AA and 10 ng/mL bFGF), and maintained in a 37°C, 5% CO_2_ incubator for further expansion.

### siRNA Transfection

Mouse primacy oligodendrocyte precursor cells (OPCs) were seeded onto PDL-coated 6-well plate at a density of 2×10^5^ cells/well a day before transfection. Cells were transiently transfected with either siRNA targeting *Prkce* or non-targeting control (Origene, SR427452) at a final concentration of 30 nM using X-tremeGENE 360 Transfection Reagent (Roche, 8724105001). After 24 hr of knockdown, cells were cultivated with either proliferating (supplemented with growth factors) or differentiation (supplemented T3, 60 ng/mL) media. After 3 days of proliferation and 5 days of differentiation, cells were harvested, and proteins were extracted and processed for western blot analysis.

### Qualitative lipidomic analysis of samples by electrospray triple Quadrupole mass spectrometry coupled with high performance liquid chromatography

Total lipids were extracted from frozen 40-70 mg human brain dissected as described above. Lipidomics profiling in mouse plasma and tissue samples was performed using Ultra Performance Liquid Chromatography-Tandem Mass Spectrometry (UPLC-MSMS). Lipid extracts were prepared from homogenized tissue samples using modified Bligh and Dyer method ^100^, spiked with appropriate internal standards, and analyzed on a platform comprising Agilent 1260 Infinity HPLC integrated to Agilent 6490A QQQ mass spectrometer controlled by Masshunter v 7.0 (Agilent Technologies, Santa Clara, CA). Glycerophospholipids and sphingolipids were separated with normal-phase HPLC as described before ^101^, with a few modifications. An Agilent Zorbax Rx-Sil column ( 2.1 x 100 mm, 1.8 µm) maintained at 25°C was used under the following conditions: mobile phase A (chloroform: methanol: ammonium hydroxide, 89.9:10:0.1, v/v) and mobile phase B (chloroform: methanol: water: ammonium hydroxide, 55:39:5.9:0.1, v/v); 95% A for 2 min, decreased linearly to 30% A over 18 min and further decreased to 25% A over 3 min, before returning to 95% over 2 min and held for 6 min. Separation of sterols and glycerolipids was carried out on a reverse phase Agilent Zorbax Eclipse XDB-C18 column (4.6 x 100 mm, 3.5um) using an isocratic mobile phase, chloroform, methanol, 0.1 M ammonium acetate (25:25:1) at a flow rate of 300 μl/min. Quantification of lipid species was accomplished using multiple reaction monitoring (MRM) transitions ^101,102^ under both positive and negative ionization modes in conjunction with referencing of appropriate internal standards: PA 14:0/14:0, PC 14:0/14:0, PE 14:0/14:0, PG 15:0/15:0, PI 17:0/20:4, PS 14:0/14:0, BMP 14:0/14:0, APG 14:0/14:0, LPC 17:0, LPE 14:0, LPI 13:0, Cer d18:1/17:0, SM d18:1/12:0, dhSM d18:0/12:0, GalCer d18:1/12:0, GluCer d18:1/12:0, LacCer d18:1/12:0, D7-cholesterol, CE 17:0, MG 17:0, 4ME 16:0 diether DG, D5-TG 16:0/18:0/16:0 (Avanti Polar Lipids, Alabaster, AL). Lipid levels for each sample were calculated by summing up the total number of moles of all lipid species measured by all three LC-MS methodologies, and then normalizing that total to mol %. The final data are presented as mean mol % with error bars showing mean ± S.E. Statistical comparisons were done using a one-way ANOVA and Tukey’s test for post-hoc analysis. Only results on DAG are provided.

### Western blots

#### Mouse

Brain tissue was prepared for western blot analysis as follows: Soluble/Insoluble Fractionation: Striatal tissue was processed as described previously ^103^. Total Fractionation: Isolated striatum or cortex was homogenized with 20 strokes of a potter-Elvenhjem glass tissue homogenizer in 1mL modified RIPA buffer (50 mM Tris-HCl pH 7.4, 1% NP-40, 0.25% Na-deoxycholate, 150 mM NaCl, 1mM EDTA) supplemented with one Pierce protease inhibitor mini tablet (Fisher Scientific A32953), 1mM PMSF, phosphatase inhibitors 2 (Millipore Sigma, P5726) (1:1000) and 3 (Millipore Sigma P0044) (1:1000), 10 μg/mL aprotinin, and 10 μg/mL leupeptin. Lysates were sonicated then centrifuged at 16,000 rcf for 15 minutes, and 5-10μg analyzed by western blot. Combined linear range was quantified on Empiria by analyzing a concentration gradient of protein (1.25, 2.5, 5, 10, and 20 μg per lane) with Revert for each antibody (Licor) to determine loading concentration. Protein was then subjected to SDS/PAGE on a NuPage Novex 4-12% Bis-Tris precast gel (Thermo Fisher NW04125) with MOPS running buffer (Invitrogen NP0001) and transferred onto a Immobilon-FL PVDF (Millipore Sigma IPFL00010) membrane. 5µg of reduced, insoluble protein from Insoluble Fractions were resolved on 3-8% Tris-Acetate Poly-Acrylamide gels. Whole protein was quantified using the revert assay (LI-COR Biosciences 926-11016), and the membrane was blocked with Intercept (TBS) Blocking Buffer (LI-COR biosciences 927-60010) for 1 hour. The membrane was then incubated in primary antibodies overnight, washed three times with TBS-0.1% Tween-20, and incubated for 1 hour in Intercept block supplemented with 0.1% Tween-20 and near-infrared conjugated secondary antibodies. Membranes were imaged on a LI-COR scanner and quantified using Empiria Software. Experiments were performed at least twice with multiple biological replicates. Antibodies for the following antigens were used DGKB (Thermofisher cat# PA5-15416 1:1000), PRKCE (Invitrogen PA5-83725 – 1:1000), p-PKCε (ser729) (Millipore 06-821-1; 1:1000), SGK1 (abcam - ab59337 1:1000), TPK1 (Fisherscientific cat# 50-172-6732 1:500), GPI1 (Thermofisher cat# PA5-26787 1:1000), Anti-Huntingtin Antibody, a.a. 1-82 | MAB5492 - EMD Millipore. The mice used for westerns were from two separate cohorts and did not include the mice used for snRNAseq and snATACseq. 6 males animals per group were used for each western except for the solb/insolb fractionated western which were mice were from a third cohort that included 4 male mice per group. All Western statistical analysis was performed using Students T-Test with two-tailed distribution and two-sample equal variance (homoscedastic). Exact p-values for significant differences are provided in the figure.

#### Human

Protein was extracted from dissected frozen tissue using RIPA buffer on ice. Protein concentration was estimated using a modified Bradford assay. Western blotting was performed using sodium dodecyl sulfate–polyacrylamide gel electrophoresis (SDS-PAGE) as described previously ^104^. Briefly, protein lysates were separated by precast 4-20 % Bis-Tris gradient gels (GenScript), followed by transferring onto PVDF membrane (Millipore). After 1 hour blocking in blocking buffer (5% milk, 0.1% TBS-Tween) at room temperature, membranes were incubated overnight at 4°C with primary antibodies. Antibodies for the following antigens were used MAG (Proteintech cat#14386-1-AP - 1:1000-3000), MOG (Proteintech #12690-1-AP - 1:500-1000), PRKCE (Invitrogen PA5-83725 – 1:1000), p-PKCε (ser729) (Millipore 06-821-1; 1:1000), MBP (Cell signal #78896S, 1:1000), SGK1 (abcam - ab59337 1:1000), TPK1 (Fisherscientific cat# 50-172-6732 1:500), GAPDH (Proteintech 60004-1-Ig 1:1000), Actin (Proteintech 66009-1-Ig; 1:5000), Anti-mouse and anti-rabbit Peroxidase-AffiniPure Donkey IgG (H+L) (Jackson ImmunoResearch Labs Cat# 715-035-151 and 711-035-152). Detection was using enhanced chemiluminescence (cat# 1705061 or 1705062) on a Bio-Rad ChemiDoc™ Touch Imaging System. Band areas were normalized to Actin and/or GAPDH. Statistical comparisons were conducted using unpaired two-tailed t-test or Mann-Whitney test as appropriate. TPK1 was analyzed separately using similar methods as described in the mouse section, and using only striatal tissue lysates.

Western blot analysis of OPC cultures was performed as outlined above with the following modifications. The following antibodies were used: rabbit anti-PRC-epsilon (Invitrogen PA5-83725, 1:1000), mouse anti-OLIG2 (Millipore, MABN50, 1:1000), mouse anti-CNPase (Biolegend, SMI-91, 1:5000), rabbit anti-MOG (Thermo, PA5-19602, 1:1000) and mouse anti-αTUBULIN (Calbiochem, CP06, 1:2500). Detection of target proteins was done by measuring chemiluminescence signal using ECL™ Prime Western Blotting Detection Reagent (Sigma, GERPN2232) on a ChemiDoc Imaging System (Bio-Rad). Image J was used to quantify the protein bands and αTUBULIN was used as loading control.

### Immunohistochemistry and in situ hybridization

Standard chromogenic and fluorescent immunohistochemistry as well as in situ hybridization were done as described previously ^94^. Paraffin-embedded formalin-fixed tissue sections were used for IHC and ISH. The following antibodies were used CA2 (Abcam ab124687- 1:100), MBP (Invitrogen PA1-10008 – 1:5000). RNAscope™ was done per the manufacturer instructions using an RNAscope ™ multiplex Fluorescent v2 kit (ACDbio 323100) with the following probes for SPP1 (cat# 889751-C2), NEAT1 (cat# 411531-C3), and MBP (cat# 573051-C4).

### Imaging and quantification

Whole slides were scanned and the images on an Aperio™ Leica slide scanner at 40X. Fluorescent stained slides were scanned on Leica Aperio™ Versa scanner at 40X. additional images were taken on a Zeiss™ 810 LSM 800 confocal microscope at using a 40X/1.3 NA oil-immersion objective. For quantification of IHC, we employed an automated method using Qupath v0.2 positive cell detection algorithm ^105^. Identification of pencil fibers and blood vessels was done using a pixel classifier trained on regions not quantified but in the same slide. Quantification of ISH slides uses positive cell detection method followed by subcellular detection. Only cells with nuclear signal were considered positive. Staining artifact and blood vessels were excluded. One or more images from each patient were used. The results were loaded in R v4.0. and cells with a minimum of 3 or more MBP dots or clusters were considered positive. NEAT1 and SPP1 were quantified in MBP positive cells. Nuclei with 2 or more dots or clusters were considered positive for SPP1 and with 2 or more dots/clusters for NEAT1. Statistical comparisons were done using one-tailed t-test or Wilcox rank test as appropriate. For calculating MBP:CA2 ratios, immunofluorescence for MBP and CA2 was performed on three or more images per case from 3 HD and 4 control caudate stained sections. The MBP signal was binarized using the threshold function in ImageJ (threshold detected automatically) and was divided by the number of CA2 positive cells counted in each image.

### Statistical analyses

All features highlighted in the paper and reported as statistically significant have p-values < 0.05 or adjusted p-values < 0.1, unless otherwise stated.

## Supporting information

Supplemental Results

## Data and code availability

All data and code are available from the corresponding authors upon reasonable request.

Data for this study can be found at:

GEO accession numbers:

Human data: GSE180928

Mouse snRNAseq: GSE180294

Mouse ATACseq: GSE180236

## Acknowledgements

We would like to thank Iliana Herrera and Marie Heath for their technical assistance, and Karen Sachs for guidance with causal network modeling. This work was supported by the following NIH grants: R35 NS116872 and P01 NS092525 (L.M.T.), R01 NS089076 (E.F. and L.M.T.). OAD was supported through funding from the Huntington Disease Society of America and the Hereditary Disease Foundation and RGL by the Hereditary Disease Foundation. This work was possible, in part, through access to the UCI Genomic High Throughput Facility Shared Resource of the Cancer Center Support Grant (CA-62203) and the Flow Cytometry Core in the Sue and Bill Gross Stem Cell Center. This research was supported by the Digital Computational Pathology Laboratory in the Department of Pathology and Cell Biology at Columbia University Irving Medical Center, and by the Biomarkers Core Laboratory at the Irving Institute for Clinical and Translational Research, home to Columbia University’s Clinical and Translational Science Award.

## Author Contributions

OAD, RGL, JW, JCR, LMT, and JEG designed and oversaw the study. Data analyses for the snRNAseq data from the mouse model and also human OPC and OLs were performed by RGL, JW, RM, AMR, VM, FP, and OAD. RM and RGL performed the WGCNA on the mouse snRNAseq data. RGL performed the causal network modeling. EM, NM and VS performed the nuclei isolation and 10x prep for the mouse snRNAseq. OAD and FK performed nuclei isolation on human samples. MA performed the ATACseq and MPG and VS performed the ATACseq data analyses. KO performed DNA extractions and assisted in data analysis. OAD, AT, and FK performed the immunohistochemistry, in situ hybridization, imaging, and image analysis on the human data. DD, HJP, and PC, performed the in vitro studies and analysis. JPV diagnosed and assessed the human postmortem HD tissue. XF assisted with human tissue collection and clinical metadata. GT performed all human proteomic analyses. SD analyzed the lipidomic data. JCR, AL, and NG handled the mice, did brain isolations, and performed Licor western and data analyses for the mouse data. AL carried out isolation of single nuclei. RGL, OAD, JW, JEG and LMT wrote the manuscript. All authors read, edited, and approved the manuscript.

## References

1 Vonsattel, J. P., Keller, C. & Del Pilar Amaya, M. Neuropathology of Huntington’s disease. Handb Clin Neurol 89, 599–618, doi:10.1016/s0072-9752(07)01256-0 (2008).

2 The Huntington’s Disease Collaborative Research Group. A novel gene containing a trinucleotide repeat that is expanded and unstable on Huntington’s disease chromosomes. Cell 72, 971–983, doi:10.1016/0092-8674(93)90585-e (1993).

3 Consortium, The Huntington’s Disease iPSC Consortium. Developmental alterations in Huntington’s disease neural cells and pharmacological rescue in cells and mice. Nat. Neurosci. 20, 648–660, doi:10.1038/nn.4532 (2017).

4 Hyeon, J. W., Kim, A. H. & Yano, H. Epigenetic regulation in Huntington’s disease. Neurochem Int, 105074, doi:10.1016/j.neuint.2021.105074 (2021).

5 Malla, B., Guo, X., Senger, G., Chasapopoulou, Z. & Yildirim, F. A Systematic Review of Transcriptional Dysregulation in Huntington’s Disease Studied by RNA Sequencing. Front Genet 12, 751033, doi:10.3389/fgene.2021.751033 (2021).

6 Vashishtha, M. et al. Targeting H3K4 trimethylation in Huntington disease. Proc. Natl. Acad. Sci. U. S. A. 110, E3027–3036, doi:10.1073/pnas.1311323110 (2013).

7 Barnat, M. et al. Huntington’s disease alters human neurodevelopment. Science 369, 787–793, doi:10.1126/science.aax3338 (2020).

8 Hickman, R. A. et al. Developmental malformations in Huntington disease: neuropathologic evidence of focal neuronal migration defects in a subset of adult brains. Acta Neuropathol 141, 399–413, doi:10.1007/s00401-021-02269-4 (2021).

9 Conforti, P. et al. Faulty neuronal determination and cell polarization are reverted by modulating HD early phenotypes. Proc Natl Acad Sci U S A 115, E762–E771, doi:10.1073/pnas.1715865115 (2018).

10 Lee, C. Y., Cantle, J. P. & Yang, X. W. Genetic manipulations of mutant huntingtin in mice: new insights into Huntington’s disease pathogenesis. FEBS J 280, 4382–4394, doi:10.1111/febs.12418 (2013).

11 Ferrari Bardile, C., et al. Intrinsic mutant HTT-mediated defects in oligodendroglia cause myelination deficits and behavioral abnormalities in Huntington disease. Proc Natl Acad Sci U S A 116, 9622–9627, doi:10.1073/pnas.1818042116 (2019).

12 Lim, R. G. et al. Huntington’s Disease iPSC-Derived Brain Microvascular Endothelial Cells Reveal WNT-Mediated Angiogenic and Blood-Brain Barrier Deficits. Cell Rep. 19, 1365–1377, doi:10.1016/j.celrep.2017.04.021 (2017).

13 Haremaki, T. et al. Self-organizing neuruloids model developmental aspects of Huntington’s disease in the ectodermal compartment. Nat Biotechnol 37, 1198–1208, doi:10.1038/s41587-019-0237-5 (2019).

14 Lee, H. et al. Cell Type-Specific Transcriptomics Reveals that Mutant Huntingtin Leads to Mitochondrial RNA Release and Neuronal Innate Immune Activation. Neuron 107, 891–908.e898, doi:10.1016/j.neuron.2020.06.021 (2020).

15 Teo, R. T. et al. Structural and molecular myelination deficits occur prior to neuronal loss in the YAC128 and BACHD models of Huntington disease. Hum Mol Genet 25, 2621–2632, doi:10.1093/hmg/ddw122 (2016).

16 Jin, J. et al. Early white matter abnormalities, progressive brain pathology and motor deficits in a novel knock-in mouse model of Huntington’s disease. Hum Mol Genet 24, 2508–2527, doi:10.1093/hmg/ddv016 (2015).

17 Huang, B. et al. Mutant huntingtin downregulates myelin regulatory factor-mediated myelin gene expression and affects mature oligodendrocytes. Neuron 85, 1212–1226, doi:10.1016/j.neuron.2015.02.026 (2015).

18 Ferrari Bardile, C., et al. Abnormal Spinal Cord Myelination due to Oligodendrocyte Dysfunction in a Model of Huntington’s Disease. Journal of Huntington’s Disease 10, 377–384, doi:10.3233/JHD-210495 (2021).

19 Hodges, A. et al. Regional and cellular gene expression changes in human Huntington’s disease brain. Hum Mol Genet 15, 965–977, doi:10.1093/hmg/ddl013 (2006).

20 Labadorf, A. et al. RNA Sequence Analysis of Human Huntington Disease Brain Reveals an Extensive Increase in Inflammatory and Developmental Gene Expression. PLoS One 10, e0143563, doi:10.1371/journal.pone.0143563 (2015).

21 Benraiss, A. et al. Cell-intrinsic glial pathology is conserved across human and murine models of Huntington’s disease. Cell reports 36, 109308, doi:10.1016/j.celrep.2021.109308 (2021).

22 Meunier, C., Merienne, N., Jolle, C., Deglon, N. & Pellerin, L. Astrocytes are key but indirect contributors to the development of the symptomatology and pathophysiology of Huntington’s disease. Glia 64, 1841–1856, doi:10.1002/glia.23022 (2016).

23 Osipovitch, M. et al. Human ESC-Derived Chimeric Mouse Models of Huntington’s Disease Reveal Cell-Intrinsic Defects in Glial Progenitor Cell Differentiation. Cell Stem Cell 24, 107–122.e107, doi:10.1016/j.stem.2018.11.010 (2019).

24 Teo, R. T. Y. et al. Impaired Remyelination in a Mouse Model of Huntington Disease. Molecular neurobiology 56, 6873–6882, doi:10.1007/s12035-019-1579-1 (2019).

25 Wilson, H., Dervenoulas, G. & Politis, M. Structural Magnetic Resonance Imaging in Huntington’s Disease. Int Rev Neurobiol 142, 335–380, doi:10.1016/bs.irn.2018.09.006 (2018).

26 Myers, R. H. et al. Decreased neuronal and increased oligodendroglial densities in Huntington’s disease caudate nucleus. J Neuropathol Exp Neurol 50, 729–742, doi:10.1097/00005072-199111000-00005 (1991).

27 Gomez-Tortosa, E. et al. Quantitative neuropathological changes in presymptomatic Huntington’s disease. Ann Neurol 49, 29–34 (2001).

28 de la Monte, S. M., Vonsattel, J. P. & Richardson, E. P., Jr. Morphometric demonstration of atrophic changes in the cerebral cortex, white matter, and neostriatum in Huntington’s disease. J Neuropathol Exp Neurol 47, 516–525, doi:10.1097/00005072-198809000-00003 (1988).

29 Gabery, S. et al. Early white matter pathology in the fornix of the limbic system in Huntington disease. Acta neuropathologica 142, 791–806, doi:10.1007/s00401-021-02362-8 (2021).

30 Mangiarini, L. et al. Exon 1 of the HD gene with an expanded CAG repeat is sufficient to cause a progressive neurological phenotype in transgenic mice. Cell 87, 493–506, doi:10.1016/s0092-8674(00)81369-0 (1996).

31 Mayr, J. A. et al. Thiamine pyrophosphokinase deficiency in encephalopathic children with defects in the pyruvate oxidation pathway. Am J Hum Genet 89, 806–812, doi:10.1016/j.ajhg.2011.11.007 (2011).

32 Zeng, W. Q. et al. Biotin-responsive basal ganglia disease maps to 2q36.3 and is due to mutations in SLC19A3. Am J Hum Genet 77, 16–26, doi:10.1086/431216 (2005).

33 Labay, V. et al. Mutations in SLC19A2 cause thiamine-responsive megaloblastic anaemia associated with diabetes mellitus and deafness. Nat Genet 22, 300–304, doi:10.1038/10372 (1999).

34 Dhir, S., Tarasenko, M., Napoli, E. & Giulivi, C. Neurological, Psychiatric, and Biochemical Aspects of Thiamine Deficiency in Children and Adults. Front Psychiatry 10, 207, doi:10.3389/fpsyt.2019.00207 (2019).

35 Eden, E., Navon, R., Steinfeld, I., Lipson, D. & Yakhini, Z. GOrilla: a tool for discovery and visualization of enriched GO terms in ranked gene lists. BMC Bioinformatics 10, 48, doi:10.1186/1471-2105-10-48 (2009).

36 Tripathi, V. et al. The nuclear-retained noncoding RNA MALAT1 regulates alternative splicing by modulating SR splicing factor phosphorylation. Mol Cell 39, 925–938, doi:10.1016/j.molcel.2010.08.011 (2010).

37 Marques, S. et al. Oligodendrocyte heterogeneity in the mouse juvenile and adult central nervous system. Science 352, 1326–1329, doi:10.1126/science.aaf6463 (2016).

38 Trapnell, C. et al. The dynamics and regulators of cell fate decisions are revealed by pseudotemporal ordering of single cells. Nat Biotechnol 32, 381–386, doi:10.1038/nbt.2859 (2014).

39 Langfelder, P. & Horvath, S. WGCNA: an R package for weighted correlation network analysis. BMC Bioinformatics 9, 559, doi:10.1186/1471-2105-9-559 (2008).

40 Beckmann, N. D. et al. Multiscale causal networks identify VGF as a key regulator of Alzheimer’s disease. Nat. Commun. 11, 3942, doi:10.1038/s41467-020-17405-z (2020).

41 Bendall, S. C. et al. Single-cell mass cytometry of differential immune and drug responses across a human hematopoietic continuum. Science 332, 687–696, doi:10.1126/science.1198704 (2011).

42 Carcamo-Orive, I. et al. Analysis of Transcriptional Variability in a Large Human iPSC Library Reveals Genetic and Non-genetic Determinants of Heterogeneity. Cell Stem Cell 20, 518–532.e519, doi:10.1016/j.stem.2016.11.005 (2017).

43 Langfelder, P. et al. Integrated genomics and proteomics define huntingtin CAG length-dependent networks in mice. Nat Neurosci 19, 623–633, doi:10.1038/nn.4256 (2016).

44 Lobo, M. K., Yeh, C. & Yang, X. W. Pivotal role of early B-cell factor 1 in development of striatonigral medium spiny neurons in the matrix compartment. J. Neurosci. Res. 86, 2134–2146, doi:10.1002/jnr.21666 (2008).

45 Kusko, R. et al. Large-scale transcriptomic analysis reveals that pridopidine reverses aberrant gene expression and activates neuroprotective pathways in the YAC128 HD mouse. Mol Neurodegener 13, 25, doi:10.1186/s13024-018-0259-3 (2018).

46 Lee, H. et al. Cell Type-Specific Transcriptomics Reveals that Mutant Huntingtin Leads to Mitochondrial RNA Release and Neuronal Innate Immune Activation. Neuron 107, 891–908 e898, doi:10.1016/j.neuron.2020.06.021 (2020).

47 Savell, K. E. et al. A dopamine-induced gene expression signature regulates neuronal function and cocaine response. Sci Adv 6, eaba4221, doi:10.1126/sciadv.aba4221 (2020).

48 Usui, N. et al. Zbtb16 regulates social cognitive behaviors and neocortical development. Transl Psychiatry 11, 242, doi:10.1038/s41398-021-01358-y (2021).

49 Hinds, L. R. et al. Dynamic glucocorticoid-dependent regulation of Sgk1 expression in oligodendrocytes of adult male rat brain by acute stress and time of day. PLoS One 12, e0175075, doi:10.1371/journal.pone.0175075 (2017).

50 Gregath, A. & Lu, Q. R. Epigenetic modifications-insight into oligodendrocyte lineage progression, regeneration, and disease. FEBS Lett 592, 1063–1078, doi:10.1002/1873-3468.12999 (2018).

51 Yu, Y. et al. Olig2 targets chromatin remodelers to enhancers to initiate oligodendrocyte differentiation. Cell 152, 248–261, doi:10.1016/j.cell.2012.12.006 (2013).

52 Qin, Q. et al. Lisa: inferring transcriptional regulators through integrative modeling of public chromatin accessibility and ChIP-seq data. Genome Biol 21, 32, doi:10.1186/s13059-020-1934-6 (2020).

53 Bentsen, M. et al. ATAC-seq footprinting unravels kinetics of transcription factor binding during zygotic genome activation. Nat Commun 11, 4267, doi:10.1038/s41467-020-18035-1 (2020).

54 Cao, J. et al. The single-cell transcriptional landscape of mammalian organogenesis. Nature 566, 496–502, doi:10.1038/s41586-019-0969-x (2019).

55 Jakel, S. et al. Altered human oligodendrocyte heterogeneity in multiple sclerosis. Nature 566, 543–547, doi:10.1038/s41586-019-0903-2 (2019).

56 Selvaraju, R. et al. Osteopontin is upregulated during in vivo demyelination and remyelination and enhances myelin formation in vitro. Mol Cell Neurosci 25, 707–721, doi:10.1016/j.mcn.2003.12.014 (2004).

57 Cammer, W. & Zhang, H. Carbonic anhydrase in distinct precursors of astrocytes and oligodendrocytes in the forebrains of neonatal and young rats. Brain Res Dev Brain Res 67, 257–263, doi:10.1016/0165-3806(92)90226-m (1992).

58 Ishii, A., Fyffe-Maricich, S. L., Furusho, M., Miller, R. H. & Bansal, R. ERK1/ERK2 MAPK Signaling is Required to Increase Myelin Thickness Independent of Oligodendrocyte Differentiation and Initiation of Myelination. The Journal of Neuroscience 32, 8855, doi:10.1523/JNEUROSCI.0137-12.2012 (2012).

59 Yildirim, F. et al. Early epigenomic and transcriptional changes reveal Elk-1 transcription factor as a therapeutic target in Huntington’s disease. Proc Natl Acad Sci U S A 116, 24840–24851, doi:10.1073/pnas.1908113116 (2019).

60 Szu, J., Wojcinski, A., Jiang, P. & Kesari, S. Impact of the Olig Family on Neurodevelopmental Disorders. Frontiers in neuroscience 15, 659601–659601, doi:10.3389/fnins.2021.659601 (2021).

61 Garcia, F. J. et al. Single-cell dissection of the human brain vasculature. Nature 603, 893–899, doi:10.1038/s41586-022-04521-7 (2022).

62 Zhang, Y. et al. An RNA-Sequencing Transcriptome and Splicing Database of Glia, Neurons, and Vascular Cells of the Cerebral Cortex. The Journal of Neuroscience 34, 11929, doi:10.1523/JNEUROSCI.1860-14.2014 (2014).

63 Hubler, Z. et al. Accumulation of 8, 9-unsaturated sterols drives oligodendrocyte formation and remyelination. Nature 560, 372–376 (2018).

64 Valenza, M. et al. Dysfunction of the cholesterol biosynthetic pathway in Huntington’s disease. The Journal of neuroscience : the official journal of the Society for Neuroscience 25, 9932–9939, doi:10.1523/JNEUROSCI.3355-05.2005 (2005).

65 Valenza, M. et al. Cholesterol biosynthesis pathway is disturbed in YAC128 mice and is modulated by huntingtin mutation. Human molecular genetics 16, 2187–2198, doi:10.1093/hmg/ddm170 (2007).

66 Valenza, M. et al. Progressive dysfunction of the cholesterol biosynthesis pathway in the R6/2 mouse model of Huntington’s disease. Neurobiology of disease 28, 133–142, doi:10.1016/j.nbd.2007.07.004 (2007).

67 Block, R. C., Dorsey, E. R., Beck, C. A., Brenna, J. T. & Shoulson, I. Altered cholesterol and fatty acid metabolism in Huntington disease. J Clin Lipidol 4, 17–23, doi:10.1016/j.jacl.2009.11.003 (2010).

68 Carroll, J. B. et al. HdhQ111 Mice Exhibit Tissue Specific Metabolite Profiles that Include Striatal Lipid Accumulation. PloS one 10, e0134465, doi:10.1371/journal.pone.0134465 (2015).

69 Kacher, R. et al. CYP46A1 gene therapy deciphers the role of brain cholesterol metabolism in Huntington’s disease. Brain 142, 2432–2450, doi:10.1093/brain/awz174 (2019).

70 Luthi-Carter, R. et al. SIRT2 inhibition achieves neuroprotection by decreasing sterol biosynthesis. Proceedings of the National Academy of Sciences of the United States of America 107, 7927–7932, doi:10.1073/pnas.1002924107 (2010).

71 Kreilaus, F., Spiro, A. S., McLean, C. A., Garner, B. & Jenner, A. M. Evidence for altered cholesterol metabolism in Huntington’s disease post mortem brain tissue. Neuropathology and applied neurobiology 42, 535–546, doi:10.1111/nan.12286 (2016).

72 Asotra, K. & Macklin, W. B. Protein kinase C activity modulates myelin gene expression in enriched oligodendrocytes. J Neurosci Res 34, 571–588, doi:10.1002/jnr.490340509 (1993).

73 Baer, A. S. et al. Myelin-mediated inhibition of oligodendrocyte precursor differentiation can be overcome by pharmacological modulation of Fyn-RhoA and protein kinase C signalling. Brain 132, 465–481, doi:10.1093/brain/awn334 (2009).

74 Baron, W., de Jonge, J. C., de Vries, H. & Hoekstra, D. Regulation of oligodendrocyte differentiation: protein kinase C activation prevents differentiation of O2A progenitor cells toward oligodendrocytes. Glia 22, 121–129, doi:10.1002/(sici)1098-1136(199802)22:2<121::aid-glia3>3.0.co;2-a (1998).

75 Damato, M. et al. Protein Kinase C Activation Drives a Differentiation Program in an Oligodendroglial Precursor Model through the Modulation of Specific Biological Networks. Int J Mol Sci 22, doi:10.3390/ijms22105245 (2021).

76 Amaral, A. I., Tavares, J. M., Sonnewald, U. & Kotter, M. R. Oligodendrocytes: Development, Physiology and Glucose Metabolism. Adv Neurobiol 13, 275–294, doi:10.1007/978-3-319-45096-4_10 (2016).

77 da Rosa, P. M. et al. High-glucose medium induces cellular differentiation and changes in metabolic functionality of oligodendroglia. Mol Biol Rep 46, 4817–4826, doi:10.1007/s11033-019-04930-4 (2019).

78 Rinholm, J. E. et al. Regulation of oligodendrocyte development and myelination by glucose and lactate. J Neurosci 31, 538–548, doi:10.1523/JNEUROSCI.3516-10.2011 (2011).

79 Yan, H. & Rivkees, S. A. Hypoglycemia influences oligodendrocyte development and myelin formation. Neuroreport 17, 55–59, doi:10.1097/01.wnr.0000192733.00535.b6 (2006).

80 DeBrosse, S. D. et al. Spectrum of neurological and survival outcomes in pyruvate dehydrogenase complex (PDC) deficiency: lack of correlation with genotype. Mol Genet Metab 107, 394–402, doi:10.1016/j.ymgme.2012.09.001 (2012).

81 Freedman, D. et al. Loss of Oligodendrocytes in Mouse Model of Pyruvate Dehydrogenase Complex Deficiency and Partial Reversal by Phenylbutyrate Treatment. Translational Neuroscience 3, 53–61 (2020).

82 Zhang, S., Lachance, B. B., Mattson, M. P. & Jia, X. Glucose metabolic crosstalk and regulation in brain function and diseases. Prog Neurobiol 204, 102089, doi:10.1016/j.pneurobio.2021.102089 (2021).

83 Marce-Grau, A., Marti-Sanchez, L., Baide-Mairena, H., Ortigoza-Escobar, J. D. & Perez-Duenas, B. Genetic defects of thiamine transport and metabolism: A review of clinical phenotypes, genetics, and functional studies. J Inherit Metab Dis 42, 581–597, doi:10.1002/jimd.12125 (2019).

84 Pico, S. et al. CPEB alteration and aberrant transcriptome-polyadenylation lead to a treatable SLC19A3 deficiency in Huntington’s disease. Sci Transl Med 13, eabe7104, doi:10.1126/scitranslmed.abe7104 (2021).

85 Stuart, T. et al. Comprehensive Integration of Single-Cell Data. Cell 177, 1888–1902 e1821, doi:10.1016/j.cell.2019.05.031 (2019).

86 Petukhov, V. et al. dropEst: pipeline for accurate estimation of molecular counts in droplet-based single-cell RNA-seq experiments. Genome Biol 19, 78, doi:10.1186/s13059-018-1449-6 (2018).

87 McCarthy, D. J., Campbell, K. R., Lun, A. T. & Wills, Q. F. Scater: pre-processing, quality control, normalization and visualization of single-cell RNA-seq data in R. Bioinformatics 33, 1179–1186, doi:10.1093/bioinformatics/btw777 (2017).

88 Yang, S. et al. Decontamination of ambient RNA in single-cell RNA-seq with DecontX. Genome Biol 21, 57, doi:10.1186/s13059-020-1950-6 (2020).

89 Hao, Y. et al. Integrated analysis of multimodal single-cell data. Cell 184, 3573–3587 e3529, doi:10.1016/j.cell.2021.04.048 (2021).

90 Hafemeister, C. & Satija, R. Normalization and variance stabilization of single-cell RNA-seq data using regularized negative binomial regression. Genome Biol 20, 296, doi:10.1186/s13059-019-1874-1 (2019).

91 Welch, J. D. et al. Single-Cell Multi-omic Integration Compares and Contrasts Features of Brain Cell Identity. Cell 177, 1873–1887 e1817, doi:10.1016/j.cell.2019.05.006 (2019).

92 Raudvere, U. et al. g:Profiler: a web server for functional enrichment analysis and conversions of gene lists (2019 update). Nucleic Acids Res 47, W191–W198, doi:10.1093/nar/gkz369 (2019).

93 Lun, A. T., McCarthy, D. J. & Marioni, J. C. A step-by-step workflow for low-level analysis of single-cell RNA-seq data with Bioconductor. F1000Res 5, 2122, doi:10.12688/f1000research.9501.2 (2016).

94 Al-Dalahmah, O. et al. Single-nucleus RNA-seq identifies Huntington disease astrocyte states. Acta Neuropathol Commun 8, 19, doi:10.1186/s40478-020-0880-6 (2020).

95 Moon, K. R. et al. Visualizing structure and transitions in high-dimensional biological data. Nat Biotechnol 37, 1482–1492, doi:10.1038/s41587-019-0336-3 (2019).

96 Corces, M. R. et al. An improved ATAC-seq protocol reduces background and enables interrogation of frozen tissues. Nat Methods 14, 959–962, doi:10.1038/nmeth.4396 (2017).

97 Smith-Geater, C. et al. Aberrant Development Corrected in Adult-Onset Huntington’s Disease iPSC-Derived Neuronal Cultures via WNT Signaling Modulation. Stem Cell Reports 14, 406–419, doi:10.1016/j.stemcr.2020.01.015 (2020).

98 Sachs, K., Perez, O., Pe’er, D., Lauffenburger, D. A. & Nolan, G. P. Causal protein-signaling networks derived from multiparameter single-cell data. Science 308, 523–529, doi:10.1126/science.1105809 (2005).

99 Emery, B. & Dugas, J. C. Purification of oligodendrocyte lineage cells from mouse cortices by immunopanning. Cold Spring Harb Protoc 2013, 854–868, doi:10.1101/pdb.prot073973 (2013).

100 Bligh, E. G. & Dyer, W. J. A rapid method of total lipid extraction and purification. Can J Biochem Physiol 37, 911–917, doi:10.1139/o59-099 (1959).

101 Chan, R. B. et al. Comparative lipidomic analysis of mouse and human brain with Alzheimer disease. J Biol Chem 287, 2678–2688, doi:10.1074/jbc.M111.274142 (2012).

102 Hsu, F. F., Turk, J., Shi, Y. & Groisman, E. A. Characterization of acylphosphatidylglycerols from Salmonella typhimurium by tandem mass spectrometry with electrospray ionization. J Am Soc Mass Spectrom 15, 1–11, doi:10.1016/j.jasms.2003.08.006 (2004).

103 Ochaba, J. et al. PIAS1 Regulates Mutant Huntingtin Accumulation and Huntington’s Disease-Associated Phenotypes In Vivo. Neuron 90, 507–520, doi:10.1016/j.neuron.2016.03.016 (2016).

104 Dansu, D. K. et al. PRMT5 Interacting Partners and Substrates in Oligodendrocyte Lineage Cells. Front Cell Neurosci 16, 820226, doi:10.3389/fncel.2022.820226 (2022).

105 Bankhead, P. et al. QuPath: Open source software for digital pathology image analysis. Sci Rep 7, 16878, doi:10.1038/s41598-017-17204-5 (2017).

