## Supplemental Results for "Single nuclei RNAseq analysis of HD mouse models and human brain reveals impaired oligodendrocyte maturation and potential role for thiamine metabolism"

##### Integration of OPCs and OL from all ages and regions

The subclustered OPCs and OLs were also integrated across all ages and regions (**Supplementary Fig 3c-f**). These data show similar clusters exist in each age and region with NT cells found mainly in clusters 0, 2, and 4 representing MOL & MFOL, OPCs, and NFOL, respectively; and R6/2 cells found in clusters 1 and 3 representing a unique MOL group and COPs, respectively (**Supplementary Fig 3c & d**). Expression of OPC and OL maturation markers can be seen in **Supplementary Fig. 3e**, which suggests increased OPC commitment in R6/2 cells (COP cells with no Pdgfra and low OL marker expression (cluster 3 and 1)) and decreased OL maturation (MOL cells with downregulated OL marker expression (cluster 1)). Pseudotime analysis of the integrated data set also showed similar results with all ages and regions showing cells along a single trajectory with 1 branch point mainly consisting of R6/2 cells (**Fig 2d and Supplementary Fig 3f**). We next analyzed R6/2 versus NT differentially expressed genes in the OPC and OL clusters which revealed similar results to our non-integrated data per age and region (**Supplementary table 2**). Showing down regulation of OL maturation genes such as Mobp, Mal, Neat1, Plp1, and Cldn11 and upregulation of genes like Smarca2. While OPCs showed upregulation of Mbp, Plp1, and Smarca2. These data suggest commitment of development in R6/2 OPCs in all ages and regions, and impaired maturation in OLs in all regions and ages. The pseudotime analysis reveals this is most significant in the striatum as more R6/2 OLs reach similar pseudotime values in the R6/2 cortex, relative to the NT (**Fig. 2d and Supplementary Fig. 3f**).

##### WGCNA and Bnets

To determine how mHTT disrupts the network structure of these modules elucidated in the NT state, we conducted module preservation analysis with R6/2 data (**Supplementary Fig. 4a & b**). Changes to the overall connectivity of the module members and in the structure of the subnetworks (node-to-node connectivity (kME), (edge weight)) would represent disruption of co-expression through mHTT pathogenic mechanisms. While all modules showed high levels of preservation in the R6/2 samples (**Supplementary Fig. 4a-d**).

<u>Other bnets:</u> Further exploration of other cell type-specific bnets revealed similar data and also a few similar hub genes including Hs6st3, Erbb4, and Meg3 in the Ex neuron bnet (**Supplementary Fig. 4c**). Recurring themes were present in each of the cell type-specific bnets including GAG/proteoglycan related genes such as

Tspan7 and Gpc5 in the astrocyte bnet (**Supplementary Fig. 4b**). When searching our Ex neuron bnet we found that GPR1, RORx, and snrnp70 seemed to have an important causal role in that specific network. We next looked at our yellow neuronal module which correlated with both MSN and Ex neurons but was anti correlated with glial cells. This network seemed to show 2 large subnetworks that separate hub genes mainly identified in the MSN versus Ex bnets, one containing Frmd4b and snrnp70 and the other containing Hs6st3, Dgkb, and Cacna2d3. Generally, the prior subnetwork contained genes related to Grp1 signaling and splicing (Tra2a and Ddx5) while the latter network contained genes related to protein glycosylation and glucose metabolism (Dgkx, Hsxstx, Galntx, Gpcx). A common link between these two pathways which seem to be playing an important role by their location in the hierarchical structure of the bnet are Neuregulin/Erbb signaling and Lingo2, both showing novel causal relationships amongst themselves and child nodes in both subnetworks. Lingo2 has been shown to regulate EGF signaling and has a role in Parkinson disease [2,3]. These data show an important role for these pathways specifically in the pathogenesis of neuronal populations in HD.

Interestingly, our MSN and OPC/OL bnets are enriched for genes associated with schizophrenia [4,5]. Hypergeometric tests were used to assess statistical significance for overrepresentation of the schizophrenia genes in the causal networks. Including Reln and Pcdh15 other genes were Nrg1/3 and Erbb4, Smarca2, PLCB1 a gene involved in diacylglycerol formation, Htr4 a glycosylated transmembrane protein involved in G protein coupled receptor serotonin signaling, as well as other genes involved in synaptic function and GPCR and calcium signaling. These data connect both metabolism to these signaling pathways and suggest coordination of these genes towards pathogenesis in HD and schizophrenia. There is an emerging role of OLs in schizophrenia pathogenesis, and the genes identified in these 2 causal networks may be relevant to both diseases.

Hub genes in the Ex bnet included *Fam19A1* and *2* and *Frmd4b* which all play a role in GRP1 signaling that regulates insulin signaling and neuronal receptor trafficking [6,7], *Rora* and *Rorb* which are nuclear receptors that regulate many biological processes including development, circadian rhythm, and glucose metabolism, as well as *Snrnp70* an essential component of the spliceosome. These data indicate an important role for these pathways in cortical cells relative to striatal. Each of these hub genes showed a larger number of NT specific outward edges indicating a loss of relationship in HD, but surprisingly most of these genes were upregulated in R6/2 mice in their corresponding cell types.

#### Human snRNAseq data

To further characterize the major gene programs that drive OL and OPC clusters, we discovered correlated gene modules using the Louvain community analysis algorithm in monocle3. The gene module expression scores are plotted in heatmap by lineage, cluster, and grade (Condition) in **Supplementary Fig. 7e,** showing that gene modules were largely specific for either OPCs or OLs, and that there are cluster and grade specific modules. Module 2 was most highly expressed in cluster 7, and the GO enrichment analysis of its genes reveal they are related to immune system and cytokine signaling. Module 10 showed highest scores across HD grades, and its genes were enriched in GO terms related to response to stress, splicing, lipid and atherosclerosis, and antigen processing and presentation. Moreover, module 19 was most highly expressed in HD grades including HDJ, and its genes were related to GTPase function. Finally, module 8 was highest in cluster 2, 3, and 6 and HD grade 3, and its genes were related to ribosomal function and translation (**Supplementary Fig. 7e & f)**. The module genes and scores by cluster, condition, and lineage are provided in **Supplementary Table 9.**

#### Validation of OL pathology in human HD and mouse data

To confirm OL gene expression abnormalities in HD, we performed WB analysis for myelin related genes MBP and MAG, which were downregulated at the RNA level in both mouse and human data, hub gene SGK1, and metabolism related genes DGKB and GPI, which are dysregulated in both the mouse and human OL and OPC data. Protein levels of MBP and MOG were not significantly altered in the HD cingulate cortex (**Supplementary Fig. 8b and c**). Conversely, protein levels of MBP (but not MAG) were increased in the caudate nucleus (**Supplementary Fig. 8b and c**). Protein levels of SGK1 were significantly decreased in the cingulate and caudate of HD brains (**Supplementary Fig. 8b and c**).

Given that MBP levels were reduced at RNA levels, we were surprised to see increased MBP protein levels in the caudate. This could be explained by an increase in OL numbers. We therefore performed immunofluorescence labeling for Carbonic Anhydrase II (CA2), expressed in OL but not OPCs[52], on caudate and cingulate of control and HD cases, and MBP in the caudate (**Supplementary Fig. 8d**). An unaltered or reduced ratio of MBP to CA2 signal in HD compared to controls would indicate a relative decrease of MBP per OL. The results show a reduced MBP:CA2 labeling ratio, suggesting that despite the overall increase in MBP protein levels, there was a general decrease in MBP when normalized to oligodendrocyte numbers (**Supplementary**

**Fig. 8e**). We confirmed the increased CA2 result using chromogenic IHC on a larger cohort, which revealed a significant increase in the proportion of CA2+ cells in the caudate and cingulate (**Supplementary Fig. 9a-d**). Moreover, as previously reported, the overall cell density in the HD caudate was increased, consistent with gliosis (**Supplementary Fig. 9c**).

For additional mouse validation, we examined the protein levels of the hub genes and glucose and lipid metabolism related genes that are potentially relevant to OL pathology, including Sgk1, Gpi1 and Dgkb*,* using quantitative western analysis (Licor) (**Supplementary Fig. 9e&f**) on striatal and cortical tissue collected from additional R6/2 and NT mice (n=6/group). A significant difference was observed for the following proteins: Sgk1 levels were lower in the cortex, Dgkb levels were lower in the striatum (**Supplementary Fig. 9a-d**.

Finally, we carried out *in situ* hybridization to examine dysregulation of HD OLs in human brain. The snRNAseq results showed that OLs from the three anatomic regions upregulated transcription of *SPP1*, increased in oligodendrocytes in the cuprizone model of demyelination[40], and *NEAT1*, increased in HD and implicated in promoting neuronal survival [41]. We performed in situ hybridization for *SPP1, NEAT1*, and *MBP* in the cingulate, caudate, nucleus accumbens (**Supplementary Fig. 9g**). Of these regions, caudate and accumbens parenchymal OLs showed increased *SPP1* expression (**Supplementary Fig. 9h&i**), and caudate parenchymal OLs showed increased *NEAT1* expression (**Supplementary Fig. 9h)**. OLs in the white matter of the nucleus accumbens and caudate did not show significant changes in *NEAT1* and *SPP1* expression (**Supplementary Fig. 9h&i)**. These results are consistent with a compensatory signature of OL in HD, whereby HD OL upregulate signals to promote survival and myelination. Furthermore, the data localizes the signature to parenchymal rather than white matter OLs.

#### Supplementary Tables

**Supplementary Table 1. Mouse DEGs per cell type**

**Supplementary Table 2. Integrated mouse OPC and OL DEGs**

**Supplementary Table 3. Mouse WGCNA & cWGCNA gene members**

**Supplementary Table 4. Causal network gene members and interactions**

**Supplementary Table 5. ATACseq Tobias differential binding results**

**Supplementary Table 6. Human sample metadata and demographics**

**Supplementary Table 7. Human cluster markers regression CAG by lineage**

**Supplementary Table 8. Human DEGs OPC and OL by region, GO, and venn analysis**

**Supplementary Table 9. Human Gene module scores by lineage grade levine clusters**

**Supplementary Table 10. R61 genotype DEGs**

**Supplementary Table 11. R61 TB v Veh DEGs**

**Supplementary Table 12. R61 TB v NT DEGs**

#### Supplementary References


1 Consortium, Huntington's disease iPSC consortium. Developmental alterations in Huntington's disease neural cells and pharmacological rescue in cells and mice. *Nat. Neurosci.* **20**, 648-660, doi:10.1038/nn.4532 (2017).

2 Belle, N. M. *et al.* TFF3 interacts with LINGO2 to regulate EGFR activation for protection against colitis and gastrointestinal helminths. *Nat. Commun.* **10**, 4408, doi:10.1038/s41467-019-12315-1 (2019).

3 Vilariño-Güell, C. *et al.* LINGO1 and LINGO2 variants are associated with essential tremor and Parkinson disease. *Neurogenetics* **11**, 401-408, doi:10.1007/s10048-010-0241-x (2010).

4 Amberger, J. S., Bocchini, C. A., Schiettecatte, F., Scott, A. F. & Hamosh, A. OMIM.org: Online Mendelian Inheritance in Man (OMIM(R)), an online catalog of human genes and genetic disorders. *Nucleic Acids Res* **43**, D789-798, doi:10.1093/nar/gku1205 (2015).

5 Ishii, T. *et al.* Modeling of the Bipolar Disorder and Schizophrenia Using Patient-Derived Induced Pluripotent Stem Cells with Copy Number Variations of 5 and. *eNeuro* **6**, doi:10.1523/ENEURO.0403-18.2019 (2019).

6 Kitano, J. *et al.* Tamalin is a scaffold protein that interacts with multiple neuronal proteins in distinct modes of protein-protein association. *J. Biol. Chem.* **278**, 14762-14768, doi:10.1074/jbc.M300184200 (2003).

7 Li, J. *et al.* Grp1 plays a key role in linking insulin signaling to glut4 recycling. *Dev. Cell* **22**, 1286-1298, doi:10.1016/j.devcel.2012.03.004 (2012).

8 Chen, E. Y. *et al.* Enrichr: interactive and collaborative HTML5 gene list enrichment analysis tool. *BMC Bioinformatics* **14**, 128, doi:10.1186/1471-2105-14-128 (2013).

9 Graybiel, A. M. The basal ganglia: learning new tricks and loving it. *Curr Opin Neurobiol* **15**, 638-644, doi:10.1016/j.conb.2005.10.006 (2005).

10 Lin, H. & Peddada, S. D. Analysis of compositions of microbiomes with bias correction. *Nat Commun* **11**, 3514, doi:10.1038/s41467-020-17041-7 (2020).

11 Lobo, M. K., Karsten, S. L., Gray, M., Geschwind, D. H. & Yang, X. W. FACS-array profiling of striatal projection neuron subtypes in juvenile and adult mouse brains. *Nat Neurosci* **9**, 443-452, doi:10.1038/nn1654 (2006).

12 Al-Dalahmah, O. *et al.* Single-nucleus RNA-seq identifies Huntington disease astrocyte states. *Acta Neuropathol Commun* **8**, 19, doi:10.1186/s40478-020-0880-6 (2020).

### Supplementary Figure 1

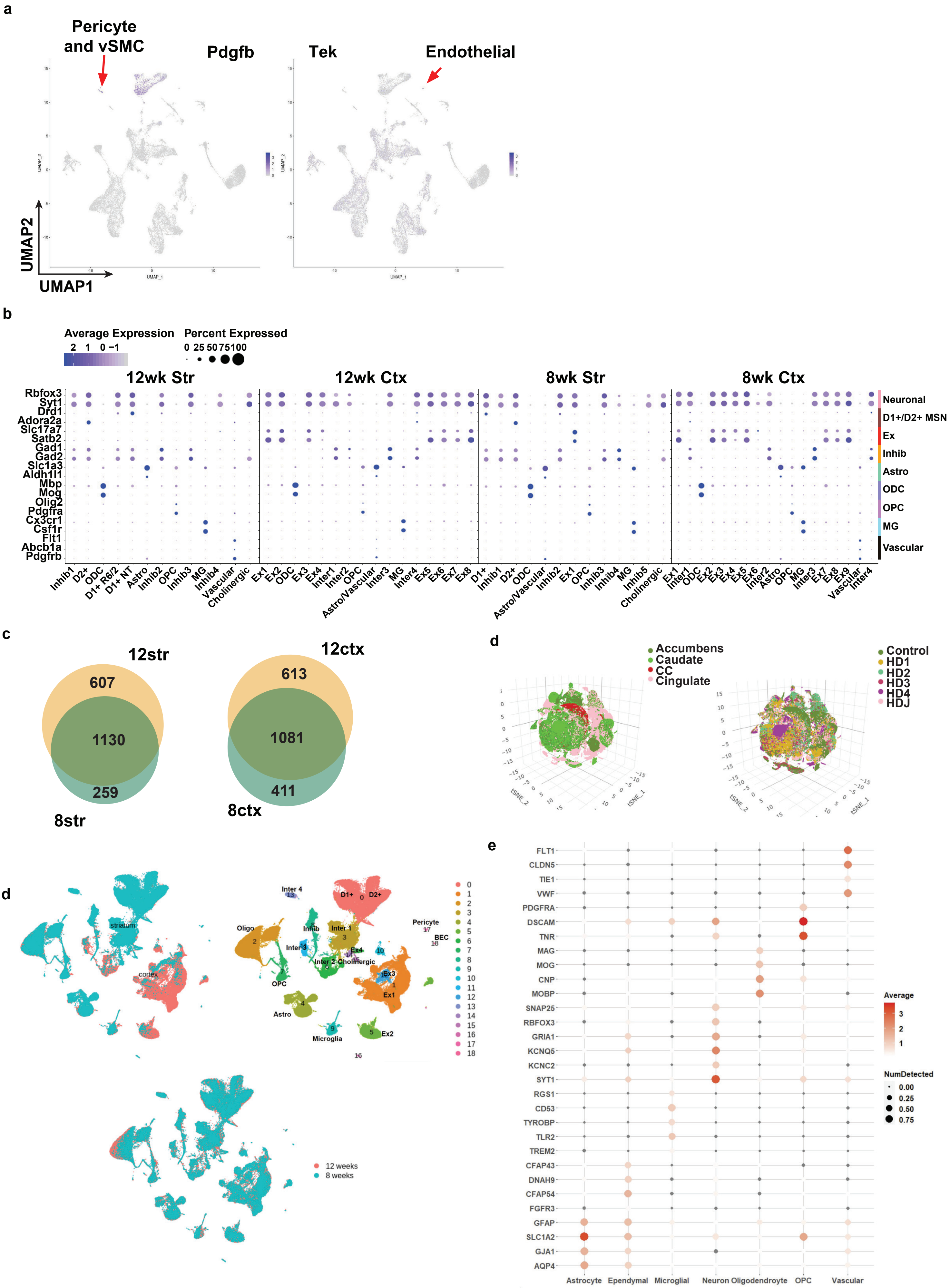

**Supplemental Figure 1. Annotation of human and mouse snRNAseq data and integrated data from both age and regions in R6/2. a**) Mouse umaps colored by expression of pdgfrb and tek showing the clustering of vascular cells with astrocytes. **b**) Dotplot showing the expression of select cell type markers across all clustered identified in the mouse data. **c**) Venn diagrams showing overlap of all DEGs between 8 and 12w, for both striatum (str) and cortex (ctx). **d)** UMAPs of integrated mouse data colored by region (top left), Cell type (top right), and age (bottom). **e)** tSNE plots of the human snRNAseq results showing color-coded by anatomic region (Top Left), and grade (Top Right). Bottom, dotplot showing expression of cell type markers per cluster.

### Supplementary Figure 2

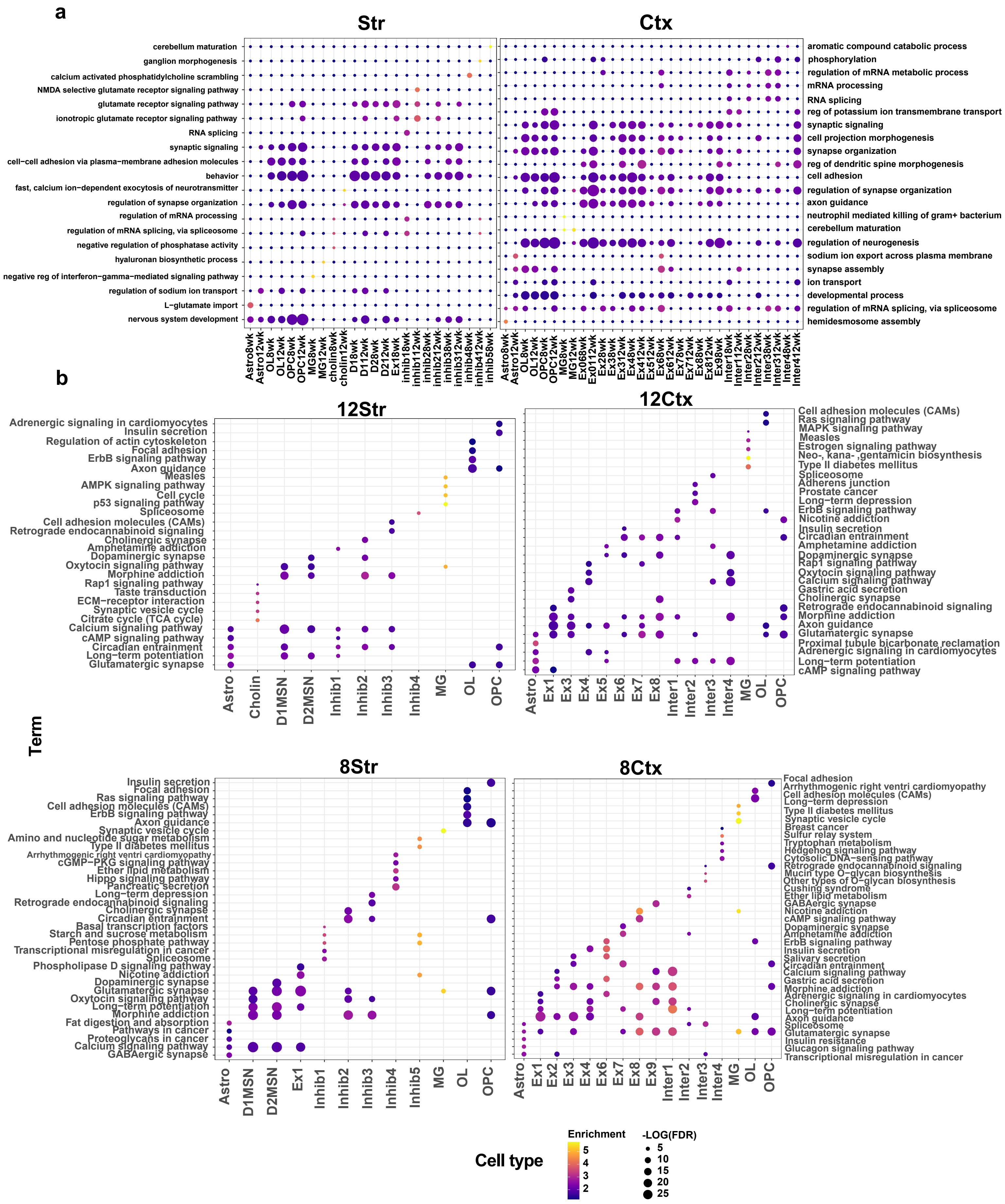

**Supplemental Figure 2. Top 5 GO terms and KEGG pathways for DEGs per a cell type and age/region.**

**a**) Top 5 GO terms per cluster in 8 and 12w striatum and cortex. **b**) Top 5 KEGG pathways in 8 and 12w striatum and cortex.

#### Supplementary Figure 3

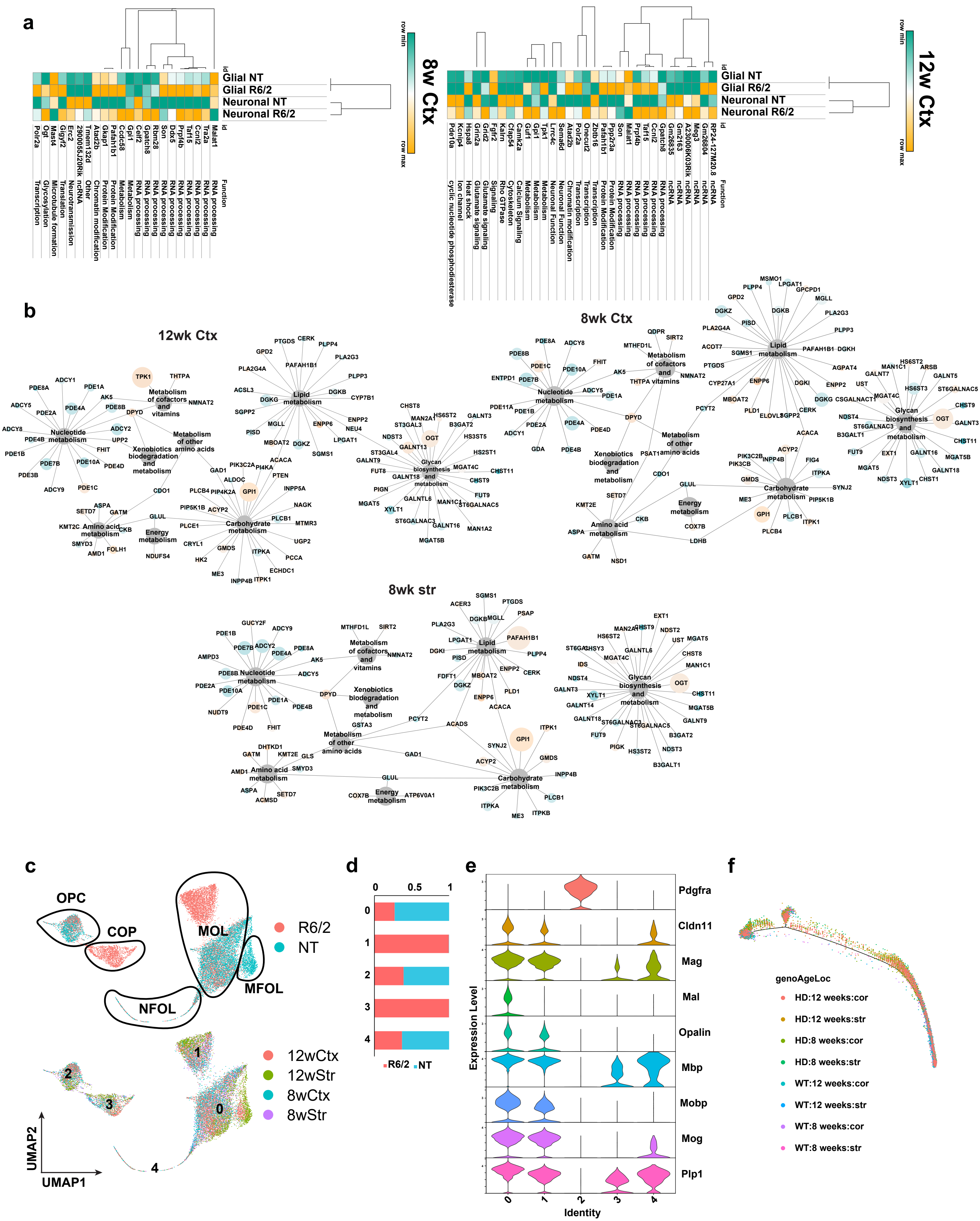

**Supplemental Figure 3. Cell type agnostic DEGs and KEGG metabolic gene networks, and integrated OPC and OL data . a**) Heatmaps and hierarchical clustering of normalized mean expression values in all glial or neuronal cells of the top cell type agnostic DEGs. Cell color represents row min (seafoam green) and max (orange). **b)** Network showing all KEGG metabolic genes significantly dysregulated across the 8wStr DEGs and both cortical dataset from every cell type. Node size is equal to the number of cell types in which the gene is found to be significantly dysregulated and node are colored by up and down regulation (orange = up and blue = down) **c**) UMAPs of integrated OPC and OL data from both ages and regions, colored by (top) genotype and (bottom) age/region. **d)** Cell number proportions by genotype in clusters 0, 1, 2, 3, and 4; corresponding to MOL, MOL, OPC, COP, NFOLs, respectively. **e**) Violin plot showing expression of OPC and OL marker and maturation genes in OPC and OL cells from all ages and regions, by cluster. **f**) Pseudotime plot of integrated OPC and OL data from both ages and regions, colored by genotype, age, and region.

#### Supplementary Figure 4

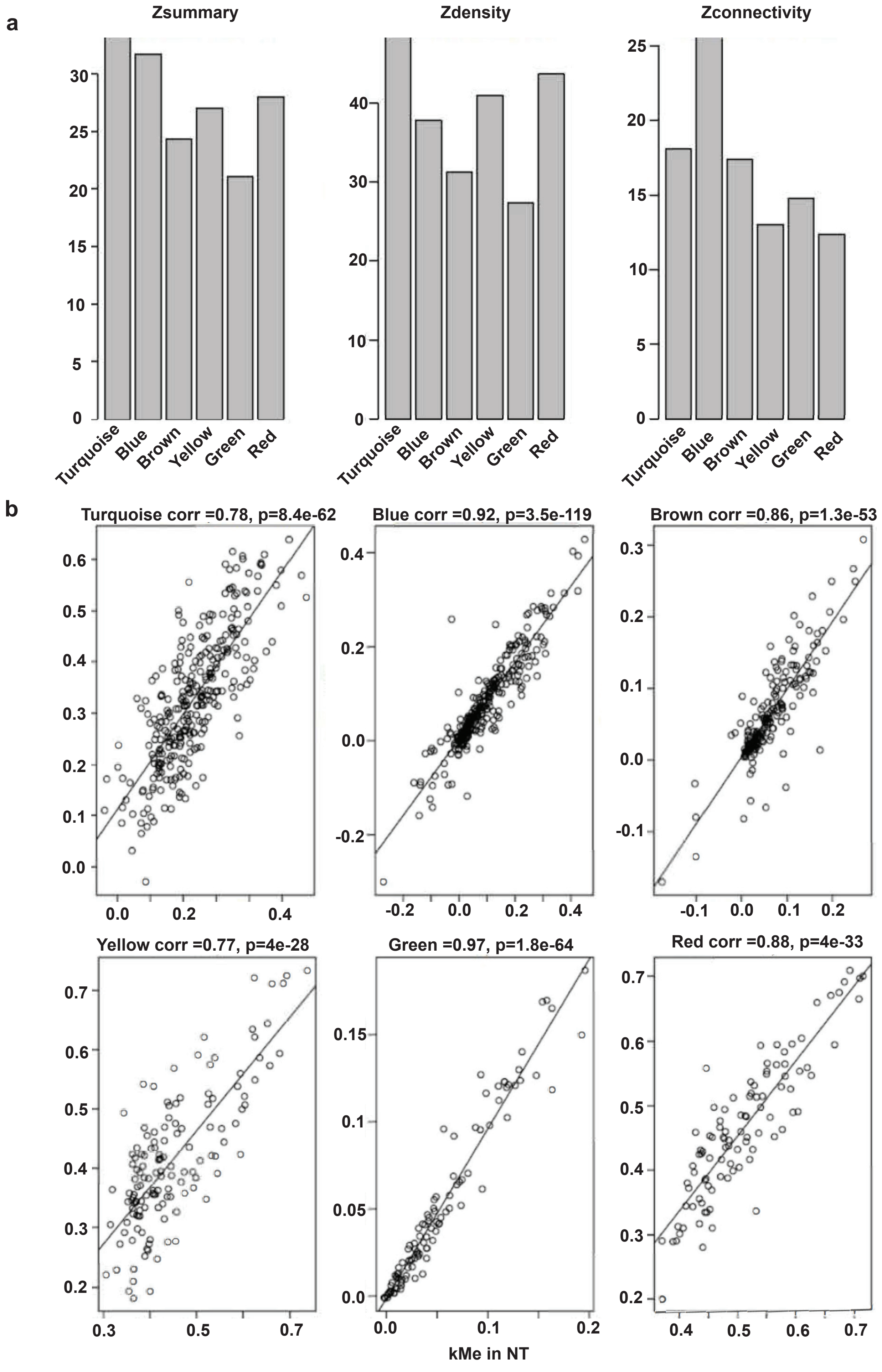

**Supplemental Figure 4. Module preservation statistics between R6/2 and NT.** Z summary preservation values > 20 for all modules and correlation > 0.78 with p values < 1.2e-53. **a**) Z-summary/density/connectivity values for module preservation. **b**) scatter plots showing kME between NT and R6/2 per module.

#### Supplementary Figure 5

a

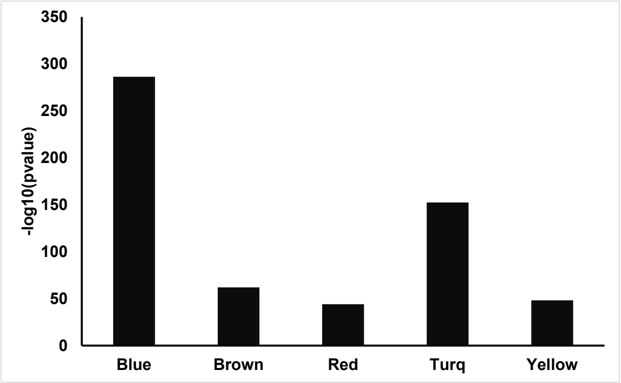


b

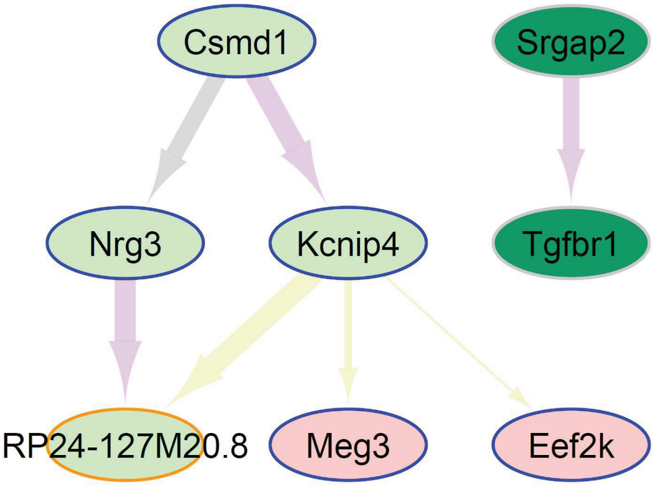


c

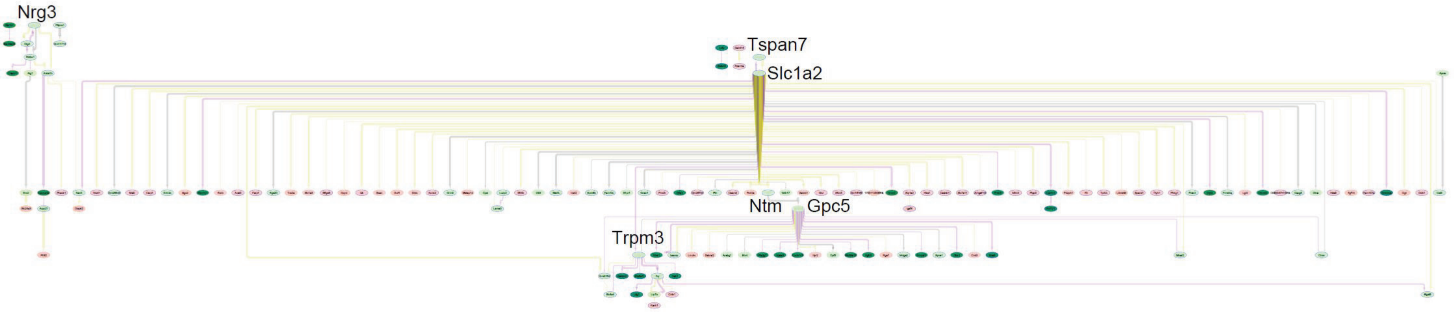


d

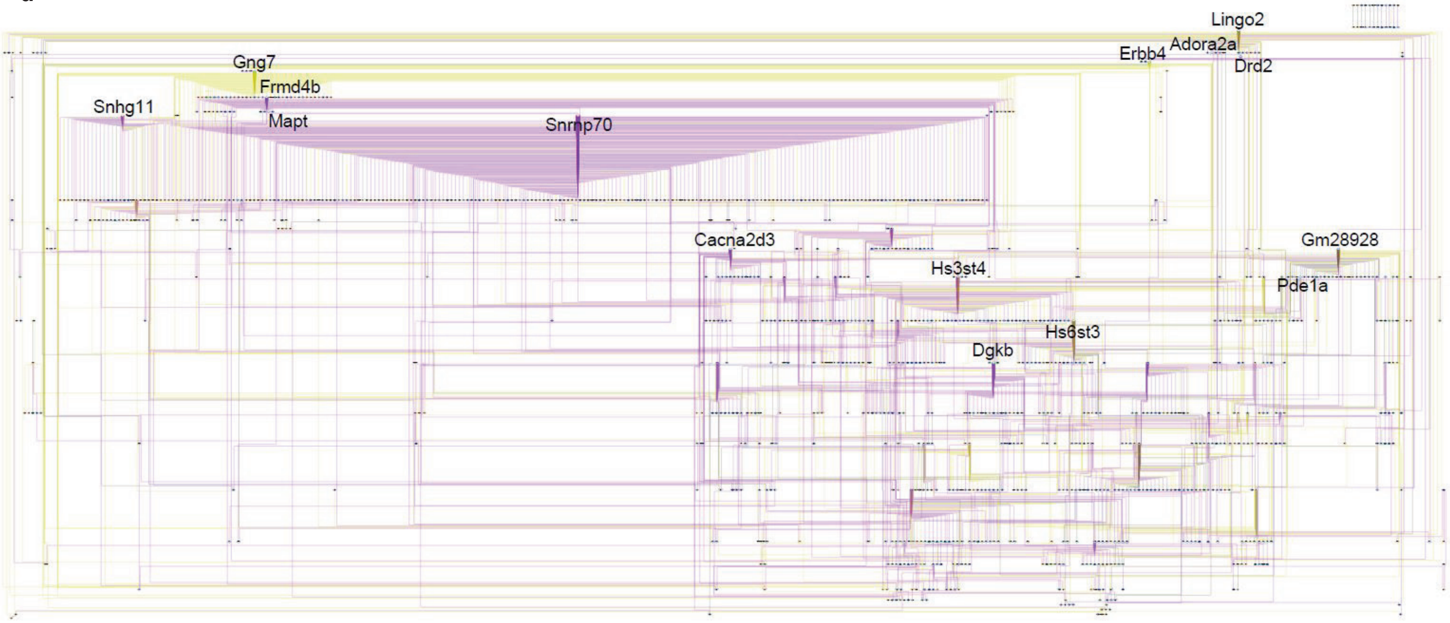

**Supplemental Figure 5. Merged causal networks for microglia, astrocyte and excitatory neurons. a**) Barplot of -log10(pvalues) from hypergeometric test of overlap between cell type DEGs and WGCNA cell type modules. **b**) Causal network for microglia. **c**) Causal network for astrocytes. **d**) Causal network for excitatory neurons. **b-d**) See **Fig. 4** legend for description of network.

#### Supplementary Figure 6

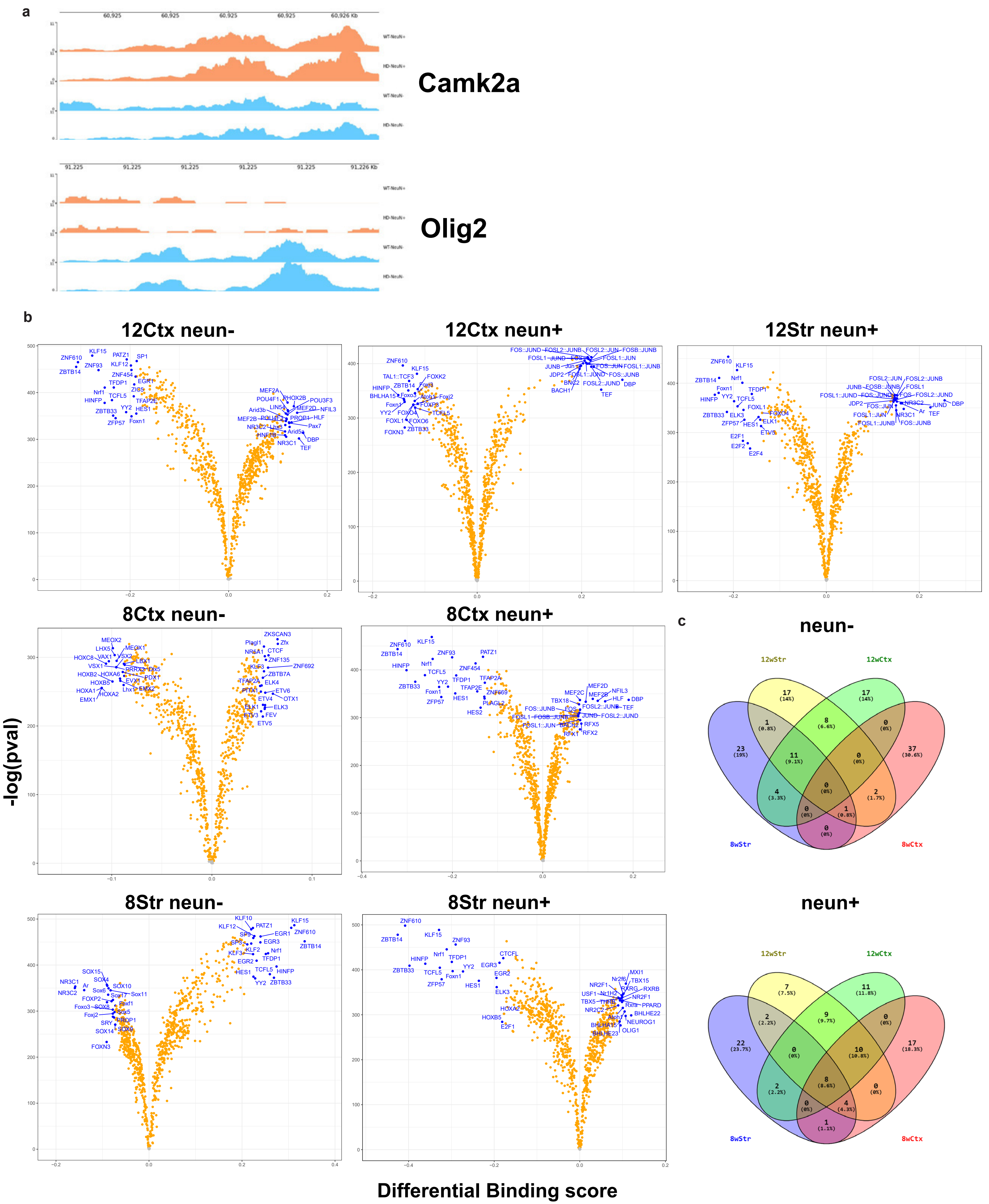

**Supplemental Figure 6. ATACSeq supplementary data. a**) Visualization of read density across camk2a and olig2 in neun+/- snRNAseq data. **b**) Volcanoplots showing differential binding scores, and -log(pvalue) differences of TF binding in open chromatin in 8 and 12w, striatum and cortex NeuN +/- cells. blue = top20 by differential binding score, orange = pvalue <0.05. **c**) Venn diagrams of overlapping TFs from ATACseq footprinting analysis per region and age.

### Supplementary Figure 7

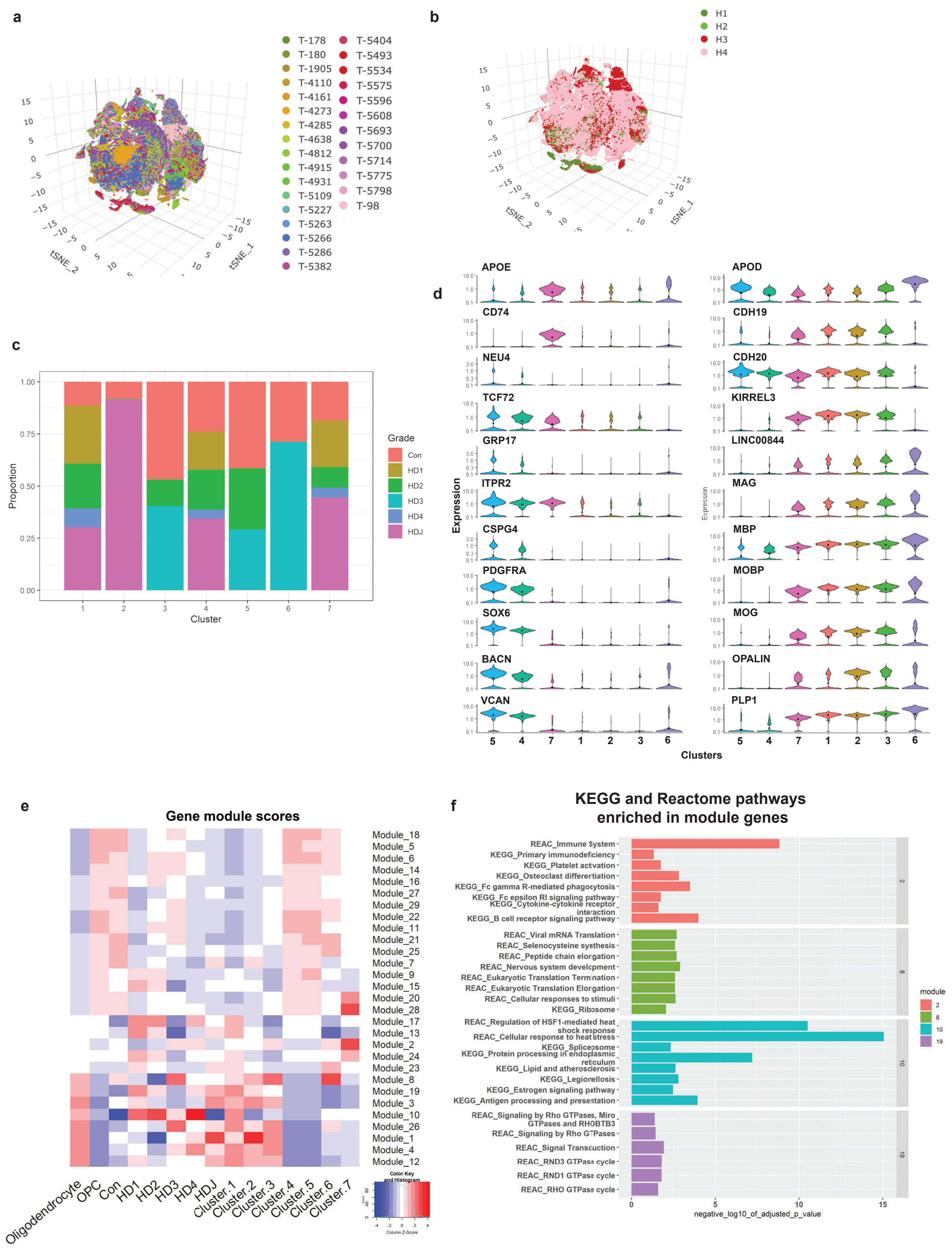

**Supplemental Figure 7. Human samples snRNAseq supplementary data. a-b**) tSNE plot showing the human snRNAseq data color-coded by donor (**a**) and sequencing batch (**b**). **c**) The relative contribution of HD grade to OL and OPC clusters is shown in bar plots. **d**) Gene expression violin plots showing the expression of select genes in OL and OPC clusters. OPC genes VCAN, BCAN, SOX6, PDGFRA, CSPG4 are most highly expressed in clusters 5 more than 4, while TCF7L2 is more expressed in cluster 4 – suggesting it is more committed. Immune OL''s genes CD74 and APOE are expressed in cluster 7. Myelin-related genes are expressed in the remaining clusters – see text for details. **e**) Gene correlation network analysis as performed in monocle3, showing the module scores against lineage, condition, and cluster. **f**) KEGG and Reactome pathway enrichment analysis in select module genes. The negative log 10 of the adjusted p value is indicated on the x-axis, and the term name on the y-axis.

#### Supplementary Figure 8

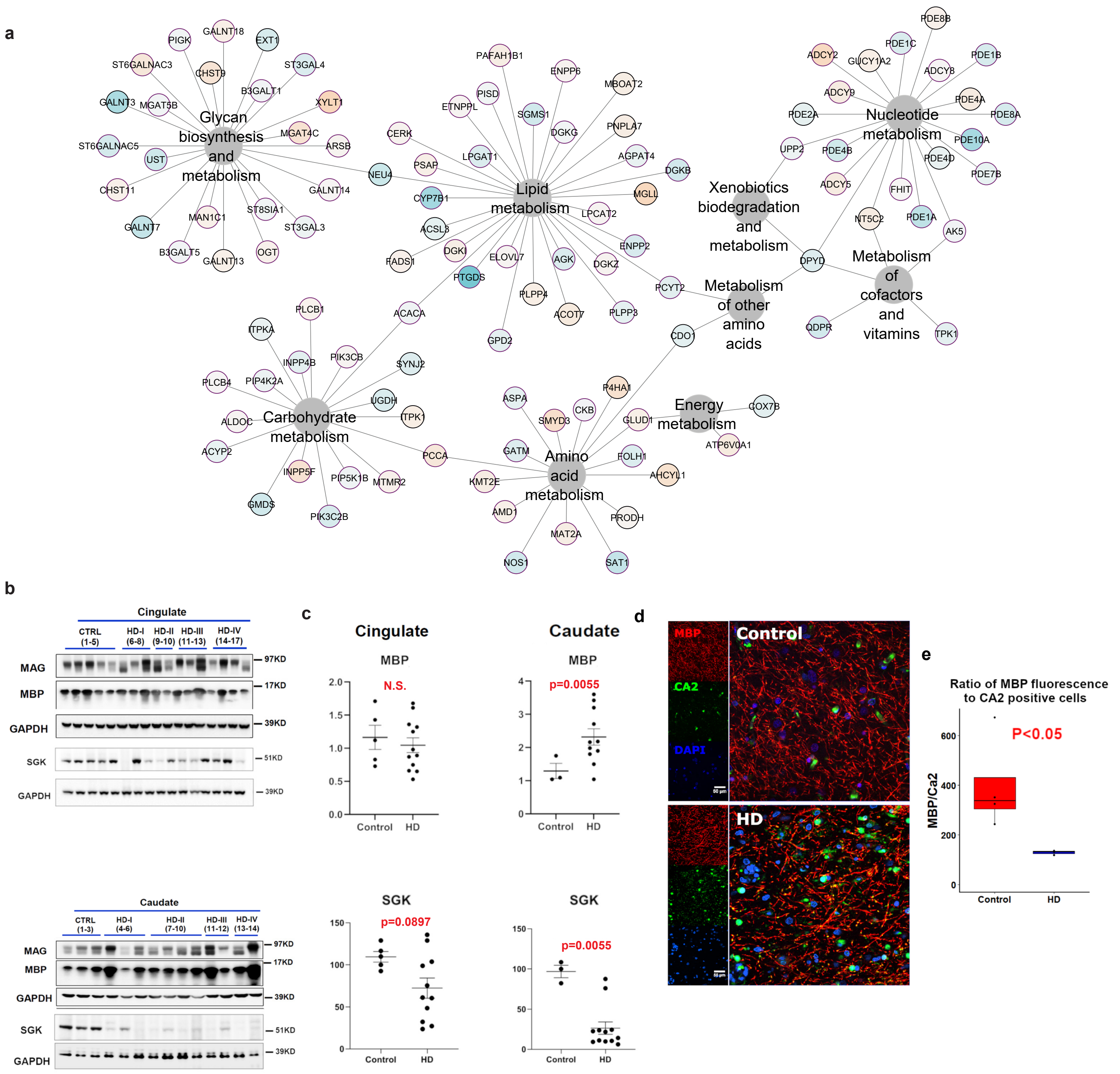

**Supplemental Figure 8. KEGG metabolic genes in human data and validation OL maturation deficits and increased OL lineage cells in the cingulate and caudate. a**) Network showing all KEGG metabolic genes significantly dysregulated across the human OPC and OL DEGs overlapping with the mouse 12w striatal DEGs. The color of the node indicates direction of DEG: orange = up and blue = down in HD. **b**) Western blot of OL maturation genes and key drivers in HD and control patient cingulate cortex and caudate. **c**) Quantification of western blow results. Mann Whitney test used for each statistical analysis. Exact p-values: Cingulate: MAG-0.2251, MBP-0.5743, SGK-0.0897; Caudate: MAG-0.2912, MBP-0.0055, SGK-0.0055. **d**) Representative images of MBP and CA2+ OLs in HD and control postmortem brain showing an increase in CA2+ OLs in the HD brain. **e**) Ratio of MBP intensity relative to CA2 positive OLs, showing a decrease in MBP per an OL. Exact p-value: 0.032.

#### Supplementary Figure 9

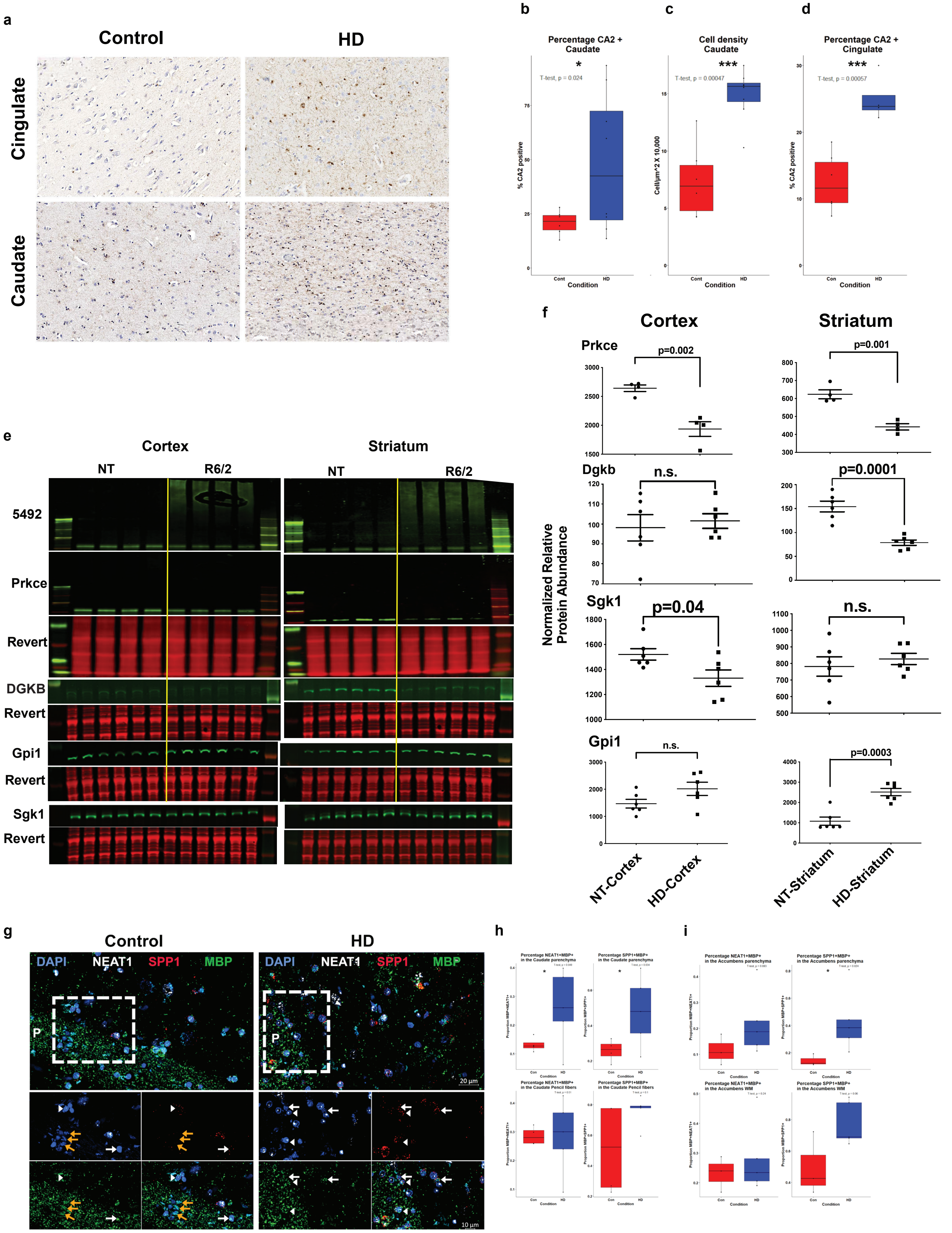

**Supplemental Figure 9. Validation of increased OL and OL stress in the caudate and accumbens, and protein validation of mouse data**. **a**) Immunohistochemical stains for Carbonic Anhydrase II (CA2), a general marker for OLs that is also expressed in early lineage OLs but not OPCs. Control and HD panels are shown in the left and right, respectively. Images of the representative regions in the cingulate cortex (Upper row) and caudate nucleus (lower row) are shown. Scale bar = 50 microns. Quantification of percentage of cells that are positive for CA2 in the caudate (**b**), and cingulate (**d**), and density of cells per unit area (**c**). The results are shown as boxplots with the bars representing the minimum and maximum values. One-tailed t-test was used to determine statistical significance. The p-values are noted on the graphs. n= 6 control and 8 for HD – for b-c, and n = 6 control and 4 HD for d. (**e and f**) Protein quantification and Licor images of select DEGs and mHTT (5492) in R6/2 and NT striatum and cortex. **e**) Licor images of mHTT (5492), Prkce in the insoluble fraction, Sgk1, Dgkb, Gpi1 and respective revert in R6/2 and NT striatum and cortex. **f**) Quantification of licor results. **g-i**) Representative images showing in situ hybridization for SPP1 (red), MBP (green), NEAT1 (white), and nuclei (DAPI - blue) in the control and HD caudate nucleus (**g**). The areas marked P represent pencil fibers of Wilson. The dashed boxes are enlarged in the lower panels. Scale bars are indicated on the graphs. Quantification of percentage of MBP- positive cells that are positive for SPP1 (right panels) and NEAT1 (left panels) in the parenchyma (upper panels) and white matter (lower panels) in the caudate (**h**), and accumbens (**i**). The results are shown as boxplots with the bars representing the minimum and maximum values. One-tailed t-test was used to determine statistical significance. The p-values are noted on the graphs. n= 4 control and 5 for HD for b, and n = 3 control and 4 HD for c.
